# Hybridization between toxic bloom-forming algae of the *Prymnesium parvum sensu lato* species complex

**DOI:** 10.64898/2026.06.03.730016

**Authors:** Nathan F. Watervoort, Timilehin Jeje, Brian P. Dilkes, Jennifer H. Wisecaver

## Abstract

The potential for hybridization to act as a driver of genetic diversity and adaptation in harmful algal bloom-forming species has received scarce attention, despite growing recognition of its occurrence in diverse protist and algal lineages. *Prymnesium parvum* s.l. (Haptophyta), is a cryptic species complex whose members form ecosystem-disruptive toxic algal blooms around the world. A prior genome analysis showed that UTEX2797, a widely used laboratory strain, originated via hybridization between clade A1 and clade A2 of this species complex. To assess the extent of A1×A2 hybridization in *P. parvum* s.l., we screened the genomes of 28 strains and identified 16 additional A1×A2 hybrid strains isolated from inland Texas or the eastern United States between 2001 and 2020. Chloroplast haplotypes indicated that hybridization between A1 and A2 may have occurred multiple times, and hybrids with different chloroplast haplotypes have been co-isolated from blooms in Texas in 2013 and 2020. Additionally, strain NIES1812 from Okinawa, Japan was sufficiently divergent from A1 and A2 to warrant designation as a separate clade, which we name A3. These results provide evidence for a facultative sexual life cycle in *P. parvum* s.l. and expand our understanding of the extensive cryptic genetic diversity present in the species complex. The frequent isolation of hybrid strains from North American blooms suggests that hybridization is common and represents a significant source of adaptive potential in these economically and ecologically damaging organisms.

## INTRODUCTION

Hybridization is the result of sexual reproduction between two genetically isolated populations or species. In plants, hybridization coupled with whole genome duplication (allopolyploidy) is a powerful evolutionary force, driving rapid diversification through the formation of new hybrid species and promoting increased rates of gene innovation and acquisition of novel traits (Soltis and Soltis 2009). Recently, whole-genome sequencing has revealed numerous examples of hybridization in fungi, including emergent pathogens (Steensels et al. 2021). In these cases, hybridization events are inferred from comparative analyses that rely on genome-level sequence data. Despite limited genomic data in most lineages of microbial eukaryotes, hybridization has been documented in ciliates (Obert et al. 2022), diatoms (Nomaguchi et al. 2018; Kim et al. 2020), dinoflagellates (Brosnahan et al. 2010), green algae (Craig et al. 2025), haptophytes (Wisecaver et al. 2023), trypanosomes (Matos et al. 2022) and other parasitic protozoa (Detwiler and Criscione 2010; Inbar et al. 2019). The growing number of hybrids identified across diverse eukaryotic lineages beyond plants and fungi suggests that hybridization may play an important evolutionary role in these groups as well.

*Prymnesium parvum* s.l. (*sensu lato*, in the broad sense) is a cryptic species complex of unicellular algae in the haptophyte lineage of eukaryotes. Species within the complex form toxic, ecosystem-disruptive algal blooms around the world (Roelke and Manning 2018; Patiño et al. 2023). *P. parvum* s.l. produce hemolytic toxins, prymnesins, that are among the largest metabolites known. The proteins responsible for prymnesin biosynthesis are comprised of ∼100 separate enzymatic domains and are encoded by massive (∼100 kbp) modular polyketide synthase genes called PKZILLAs (Fallon et al. 2024). To date, three distinct prymnesins (A-, B-, and C-type) have been identified that differ in the number of carbons in their backbone molecule, with A-type prymnesins being the largest (90 carbons) and C-types being the smallest (83 carbons) (Rasmussen et al. 2016). Relative toxicity also varies between prymnesin types with A-type prymnesins having the highest level of cytotoxicity to epithelial fish gill and human colon cell lines and B-type prymnesins having the lowest level of cytotoxicity (Varga et al. 2024). Production of the different prymnesins segregates phylogenetically across the *P. parvum* s.l. species tree forming three large A-, B-, and C-type prymnesin-producing clades (Binzer et al. 2019).

Extremely high levels of genetic diversity exist both within and among the A-, B-, and C-type clades. Haploid genome size estimates across the species complex range from 115 Mbp to 359 Mbp, and vegetatively haploid, diploid, triploid, and tetraploid strains can be maintained in stable culture (Wisecaver et al. 2023; Kuhl et al. 2024). Although cryptic speciation is recognized within *P. parvum* s.l., the boundaries between species remain unclear. Increased divergence between some A- and C-type strains suggests the presence of distinct populations or potentially multiple species within these two clades (Wisecaver et al. 2023). The habitat range of *P. parvum s.l.* appears to be expanding, and blooms are predicted to increase both in frequency and intensity in the future (Tábora-Sarmiento et al. 2022; Macêdo et al. 2023; Sobieraj and Metelski 2023). Consequently, increasing attention is being paid toward the genetic diversity present in *P. parvum s.l.* to identify genes and traits responsible for their ecological success.

Recently, Wisecaver et al. (2023) provided genome-scale evidence of hybridization within *P. parvum* s.l. clade A. A chromosome-scale de novo assembly of *P. parvum* strain UTEX2797 revealed two co-linear genomes with divergent haplotypes. One UTEX2797 haplotype grouped phylogenetically with two strains from the United States: strain 12B1 and CCMP3037, hereafter referred to as clade A1. The second UTEX2797 haplotype grouped with two strains from Europe: RCC3703 and CCMP2941, hereafter referred to as clade A2. UTEX2797 contained roughly twice as much DNA content per nucleus (∼280 Mbp, as estimated via flow cytometry), than its k-mer-based estimate of genome size (∼120 Mbp) indicating the strain was an allodiploid hybrid (Wisecaver et al. 2023). UTEX2797 was first isolated in 2002 from a Colorado River sample from Texas (United States) and has since been used in numerous growth and sequencing experiments of *P. parvum s.l.* (e.g., Blossom et al. 2014; Talarski et al. 2016; Rashel and Patiño 2017; Lysgaard et al. 2018; Anestis et al. 2021; Richardson and Patiño 2021; Bannon et al. 2024; Chávez Montes et al. 2024; Varga et al. 2024). In their analysis, Wisecaver et al. (2023) suspected that an additional strain, 12A1, also from Texas (Table 1), was likely a hybrid due to elevated rates of heterozygosity in genome sequence data comparable to that of UTEX2797. To date, the hybrid nature of strain 12A1 has not been confirmed and the extent of hybridization in *P. parvum s.l.* is unknown.

**Table 1:** Description of *P. parvum* clade A strains.

| Subclade | Strain ID | Date isolated | Isolation location | DNA content per nuclei (Mbp) | Ploidy | Dominant AIM ancestry |
| --- | --- | --- | --- | --- | --- | --- |
| **A1** | 12B1 | 2010 | Moss Creek Lake, TX, USA | 107 | haploid | A1 |
| **A1** | CC2192B4 | 2019 | Lake Colorado City, TX, USA | 271 | diploid | A1 |
| **A1** | CC2192B5 | 2019 | Lake Colorado City, TX, USA | 265 | diploid | A1 |
| **A1** | CC2192C1 | 2019 | Lake Colorado City, TX, USA | 277 | diploid | A1 |
| **A1** | CC2192D2 | 2019 | Lake Colorado City, TX, USA | 261 | diploid | A1 |
| **A1** | CC2192D3 | 2019 | Lake Colorado City, TX, USA | 261 | diploid | A1 |
| **A1** | CCMP3037 | 2008 | Twin Buttes Lake, Laramie, WY, USA | 246 | diploid | A1 |
| **A1** | PK2298B4 | 2020 | Possum Kingdom Lake, TX, USA | 120 | haploid | A1 |
| **A2** | CCMP2941 | 2007 | Repnoe, Tambov Oblast, Russia | 485* | tetraploid* | A2 |
| **A2** | RCC3703 | 1953 | Millport, Great Cumbrae, UK | 297 | diploid | A2 |
| **A3** | NIES1812 | 2005 | Yufu Isl., Okinawa, Japan | 120 | haploid | --† |
| **A hybrid** | 8020 | 2013 | Possum Kingdom Lake, TX, USA | 288 | allodiploid | A1×A2 |
| **A hybrid** | 8027 | 2013 | Possum Kingdom Lake, TX, USA | 272 | allodiploid | A1×A2 |
| **A hybrid** | 12A1 | 2010 | Moss Creek Lake, TX, USA | 275 | allodiploid | A1×A2 |
| **A hybrid** | 13A5 | 2010 | E.V. Spence Reservoir, TX, USA | 282 | allodiploid | A1×A2 |
| **A hybrid** | ARC66 | 2001 | Hilton Head, SC, USA | 270 | allodiploid | A1×A2 |
| **A hybrid** | ARC81 | 2002 | Artesian Aquafer, Elizabeth City, NC, USA | 268 | allodiploid | A1×A2 |
| **A hybrid** | ARC82 | 2002 | Artesian Aquafer, Elizabeth City, NC, USA | 283 | allodiploid | A1×A2 |
| **A hybrid** | ARC83 | 2002 | Artesian Aquafer, Elizabeth City, NC, USA | 270 | allodiploid | A1×A2 |
| **A hybrid** | ARC84 | 2002 | Artesian Aquafer, Elizabeth City, NC, USA | 273 | allodiploid | A1×A2 |
| **A hybrid** | PK2297D4 | 2020 | Possum Kingdom Lake, TX, USA | 284 | allodiploid | A1×A2 |
| **A hybrid** | PK2298C6 | 2020 | Possum Kingdom Lake, TX, USA | 280 | allodiploid | A1×A2 |
| **A hybrid** | PK2299B3 | 2020 | Possum Kingdom Lake, TX, USA | 279 | allodiploid | A1×A2 |
| **A hybrid** | PK2300D3 | 2020 | Possum Kingdom Lake, TX, USA | 269 | allodiploid | A1×A2 |
| **A hybrid** | UTEX2797 | 2002 | US Hwy 183, Texas Colorado River, TX, USA | 274 | allodiploid | A1×A2 |
| **A hybrid** | UTEX2827 | 2002 | Charleston, SC, USA | 290 | allodiploid | A1×A2 |

\* Genome size and C-value estimates for CCMP2941 are from (Wisecaver et al. 2023).
† As an outgroup, A3 strain NIES1821 showed equal proportions of homozygous A1 and A2 AIMs (see Figure 2C).

To investigate whether additional hybrid strains exist beyond UTEX2797, we screened a collection of 38 *P. parvum s.l.* strains for signatures of A1×A2 hybridization (Table S1). This included 25 *P. parvum* s.l. strains from culture collections and 10 strains isolated from recent Texas blooms. We identified numerous hybrid strains, all from North American blooms in the 21^st^ century, suggesting that hybrids are common in, and well adapted to, these environments. We present evidence that A1×A2 hybridization has likely occurred multiple times. The genomes of hybrid strains contain numerous structural variants relative to parental strains. Together, these results demonstrate that the hybrid genome found in UTEX2797 is not unusual for *P. parvum s.l.* and suggest that hybridization is an ongoing process in the evolution of *P. parvum*clade A.

## RESULTS

An analysis of SNP variation from short-read sequencing data identified 27 strains belonging to *P. parvum* clade A in the Wisecaver Lab culture collection (Table 1). In a phylogenetic network constructed from these data, strains known to produce A-, B-, and C-type prymnesins formed clear monophyletic groups, with one exception. Strain ARC140 was previously characterized as producing B-type prymnesin (Binzer et al. 2019); however, the strain grouped within the A-type clade in our analysis (Figure S1). To avoid confusion, ARC140 was excluded from downstream analyses. The remaining 26 strains that grouped within the A-type clade consisted of eight chemotyped strains with confirmed A-type prymnesin production (12A1, 12B1, ARC83, CCMP2941, CCMP3037, NIES1812, RCC3707, and UTEX2797; Binzer et al. 2019; Wisecaver et al. 2023) and 18 additional strains (Table 1).

### Variation in heterozygosity and ploidy in *P. parvum* clade A

The ploidy of all *P. parvum* clade A strains was inferred using their total nuclear DNA content compared to that of the haploid reference strain, 12B1, which has a nuclear DNA content of 107 Mbp (Table 1; Table S2). In addition to 12B1, two other strains appear haploid: NIES1812 isolated in 2005 from Okinawa, Japan (DNA content = 120 Mbp) and PK2298B4, a newly isolated strain from a 2020 bloom in Possum Kingdom Lake, Texas (DNA content = 120 Mbp) (Figure 1, Table 1). Twenty-two strains were diploid with nuclear DNA content roughly twice that of the haploid strains (average DNA content = 298 Mbp) (Figure 1, Table 1). Diploids include four strains (CCMP3037, RCC3707, UTEX2797, and 12A1) previously identified by Wisecaver et al. (2023) as well as 18 newly characterized diploid strains all isolated from the United States between 2002 and 2020, including twelve strains from Texas, four from North Carolina, and two from South Carolina (Table 1). We did not have a sample of CCMP2941 in the lab for flow cytometry at this time, but previously reported a nuclear DNA content of 485 Mbp for this strain (Wisecaver et al. 2023), indicating it is a tetraploid (Table 1).

**Figure 1:**
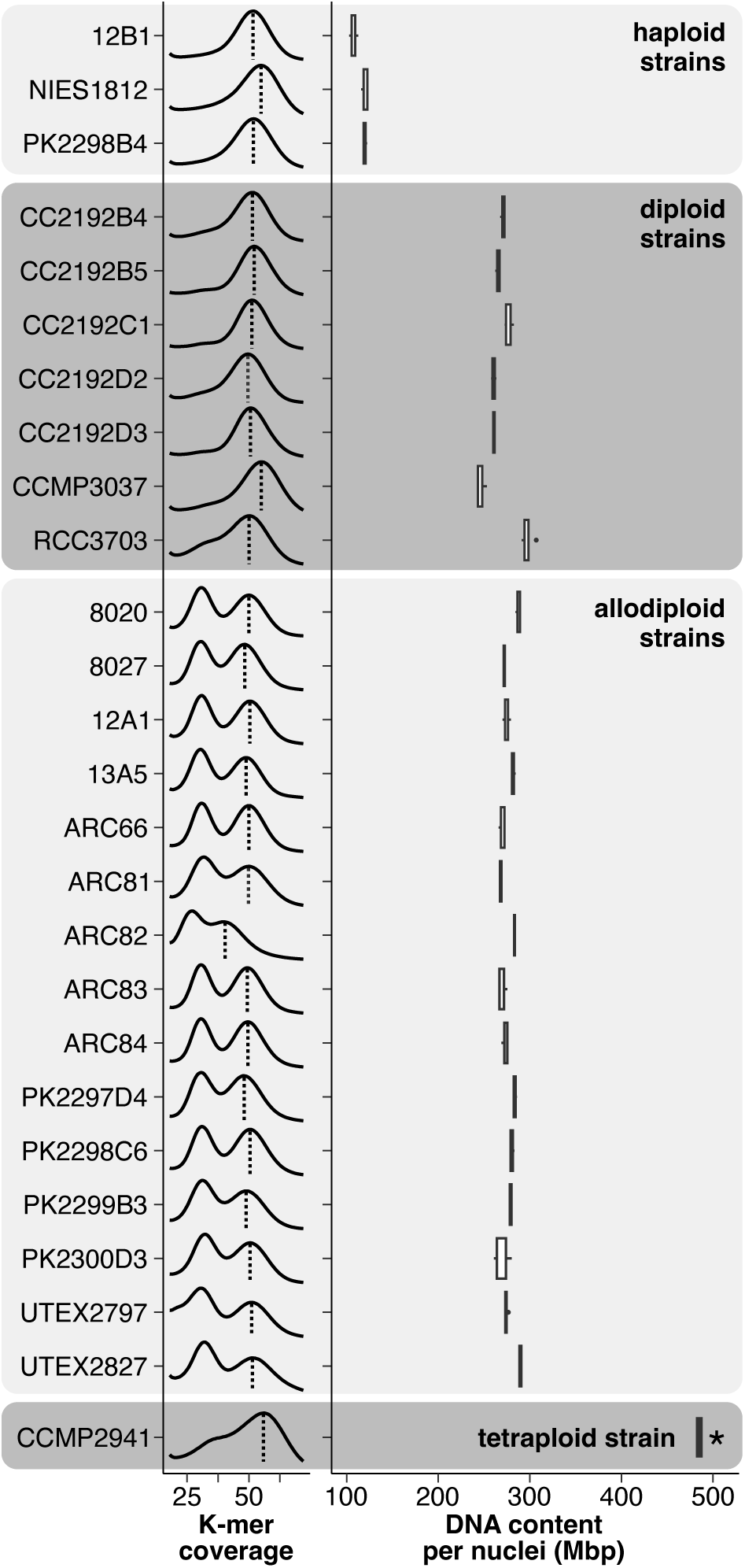
Heterozygosity and genome size variation among *P. parvum* clade A strains. Filtered Illumina reads were randomly subsampled to 10 million pairs per strain prior calculation of K-mer coverage to control for differences in sequencing depth between strains. Homozygous k-mer peaks are indicated by the vertical dashed lines. Boxplots depict total DNA content per nuclei (Mbp) based on flow cytometry. (*) Flow cytometry measurements for CCMP2941 were were taken from Wisecaver et al. (2023).

Heterozygosity was qualitatively assessed in K-mer coverage plots by the presence and relative height of a second peak at half the coverage of the homozygous peak (Figure 1, Figure S2). High heterozygosity was previously observed in strain UTEX2797 and is indicative of its hybrid origin (Wisecaver et al. 2023). Fourteen additional strains showed a high level of heterozygosity, comparable to that of UTEX2797. All 15 high-heterozygosity strains were diploid, which we classified as putative allodiploid hybrids (Figure 1).

### Phylogenetic analyses identify a new A3 lineage and additional A1×A2 hybrids

Short reads from all A-type strains were aligned to the 12B1 v1.1 nuclear genome assembly (Fallon et al. 2024) and used to call 453,986 single nucleotide polymorphisms (SNPs) shared by the genomes of at least two strains. Strains formed four clusters in a SNP-based phylogenetic network (Figure 2A). Three clusters correspond to the A1, A2, and A-hybrid subclades identified in Wisecaver et al. (2023). The A1 subclade was comprised of both haploid (12B1 and PK2298B4) and diploid strains (CCMP3037 from Wyoming and five new diploid strains isolated from a bloom in Lake Colorado City, Texas in 2019) (Figure 2A, Table 1). No additional A2 strains were identified in this analysis beyond the two already described in Wisecaver et al. (2023): RCC3703, a diploid strain from Great Cumbrae, United Kingdom, and CCMP2941, a tetraploid strain from Repnoe, Russia (Figure 2A, Table 1). All putative allodiploid hybrids identified based on K-mer frequencies and flow cytometry grouped with UTEX2797 in the A-hybrid clade (Figure 2A, Table 1). To investigate whether the hybridization signal could be due to culturing issues (e.g., two divergent diploid strains present in a single culture), we created seven single-cell reisolates of UTEX2797 (n=4) and 12A1 (n=3). These new isolates all showed the same level of heterozygosity as their progenitor cultures and grouped within the A-hybrid clade (Figure 2A, Figure S2).

**Figure 2:**
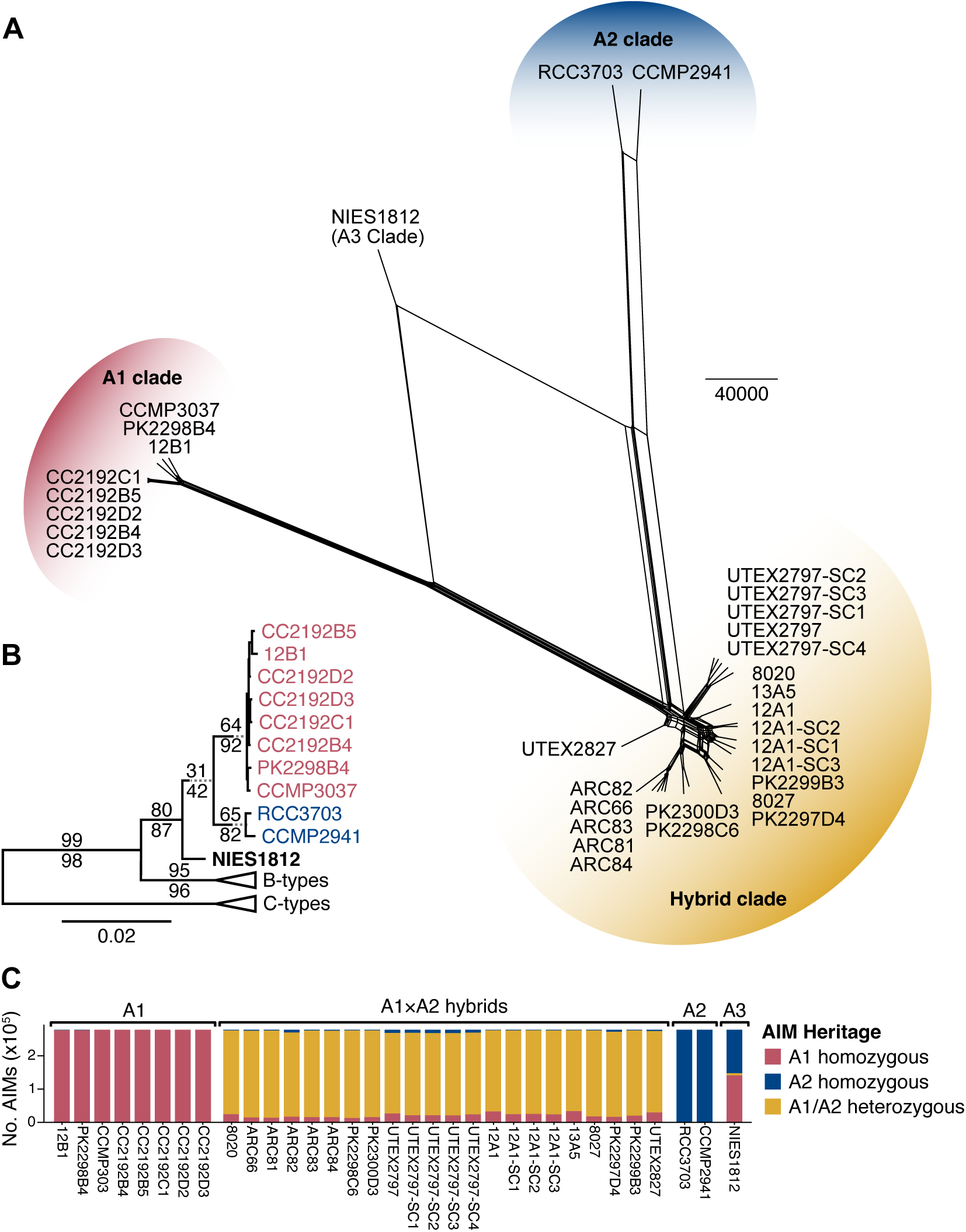
Phylogenetic analysis of *P. parvum* clade A strains. A) Network of clade A strains based on distance matrix of 453,986 SNPs. Edge lengths represent absolute SNP differences with the scale bar corresponding to 40,000 SNPs. B) Concatenation-based Maximum Likelihood species phylogeny (excluding hybrid strains) based on 9,769 single copy genes showing the relative branching pattern of clade A1 (pink), A2 (blue), and the new A3 clade comprised of NIES1812 (bolded). Support values for nodes are reported on the preceding branch as site concordance factors (above) and gene concordance factors (below). The B and C clades have been collapsed for clarify; see Figure S3 for full tree. C) Proportion of AIMs that are A1 homozygous, A2 homozygous, or A1/A2 heterozygous per strain. .

Strain NIES1812 from Okinawa, Japan grouped separately from all other strains in our analysis (Figure 2A). An earlier rDNA phylogeny placed NIES1812 as an outgroup to the A1 and A2 subclades (Binzer et al. 2019). Our SNP network was not informative with respect to the relative branching pattern of the A-type subclades as it lacked an outgroup. Therefore, to evaluate the placement of NIES1812 relative to A1 and A2, we used 9,769 single-copy genes for concatenation- and coalescent-based species tree analyses. The same species tree topology was recovered in both analyses (Figure S3) and placed NIES1812 as the outgroup to all other A-type clades in our analysis (Figure 2B). Site concordance factors (sCF) and gene concordance factors (gCF) indicated strong support for the A1 and A2 subclades as well as the B- and C-type clades (Figure 2B). However, the analysis indicated a moderate amount of discordance regarding the placement of NIES1812 (sCF=31, gCF=42, Figure 2B) relative to the A1 and A2 subclades.

Of the 453,986 SNPs in our analysis, 61.22% (n=277,926) were completely fixed for different alleles in diploid A1 strains compared to A2 strains. At these fixed sites, hereafter referred to as Ancestrally Informative Markers (AIMs), allodiploid hybrids are expected to be fully heterozygous, provided several assumptions. These assumptions are 1) that the A1 and A2 subclades represent the true parental lineages of hybrids; 2) that the SNPs are truly fixed between the A1 and A2 lineages; and 3) that minimal subsequent backcrossing or genome evolution has taken place in hybrid strains. Matching this expectation, the majority of AIMs in all allodiploid hybrids strains were heterozygous (Figure 2C; Table S3), lending further support to these strains being A1×A2 hybrids. On the high end, 94.68% of AIMs were heterozygous in strain PK2298C6, 4.72% were homozygous for the A1 allele, and 0.60% were homozygous for the A2 allele (Figure 2C; Table S3). The hybrid strain with the lowest level of heterozygosity was UTEX2797; 87.23% of AIMs were heterozygous, 9.55% were homozygous for the A1 allele, and 3.22% were homozygous for the A2 allele (Figure 2C; Table S3). In contrast to the pattern observed in hybrid strains, AIMs in NIES1812 were predominantly homozygous (as expected for a haploid strain) for either the A1 allele (50.77%) or the A2 allele (47.14%) (Figure 2C; Table S3). These SNPs did not show evidence of A3 arising as a homoploid hybrid of A1 and A2 as AIMs were evenly distributed across the genome (Figure S4). This pattern is consistent with its phylogenetic placement sister to A1/A2 and the AIM alleles present in NIES1812 likely representing the ancestral state present in their common ancestor.

### A1×A2 hybrids contain either A1- or A2-like chloroplasts and display complex evolutionary relationships

To investigate chloroplast inheritance patterns in A1×A2 hybrids, short reads from A1, A2, and A1×A2 hybrid strains were aligned to the 12B1 v1 chloroplast genome assembly (Wisecaver et al. 2023) and used to identity 188 chloroplast-encoded SNPs. Chloroplast SNPs were strongly linked and formed two distinct chloroplast haplotypes (cpHaps) corresponding to the A1 and A2 subclades (Figure 3A). Consistent with single-strain cultures of hybrids, no evidence of chloroplast heteroplasmy was detected in hybrid strains. Instead, nine hybrid strains carried an A1-like chloroplast (hereafter referred to as cpHap-A1), and six hybrids carried an A2-like chloroplast (hereafter referred to as cpHap-A2; Figure 3A). All single-cell reisolates of UTEX2797 (cpHap-A1) and 12A1 (cpHap-A2) had the same cpHap as their progenitor cultures (Figure 3A).

**Figure 3:**
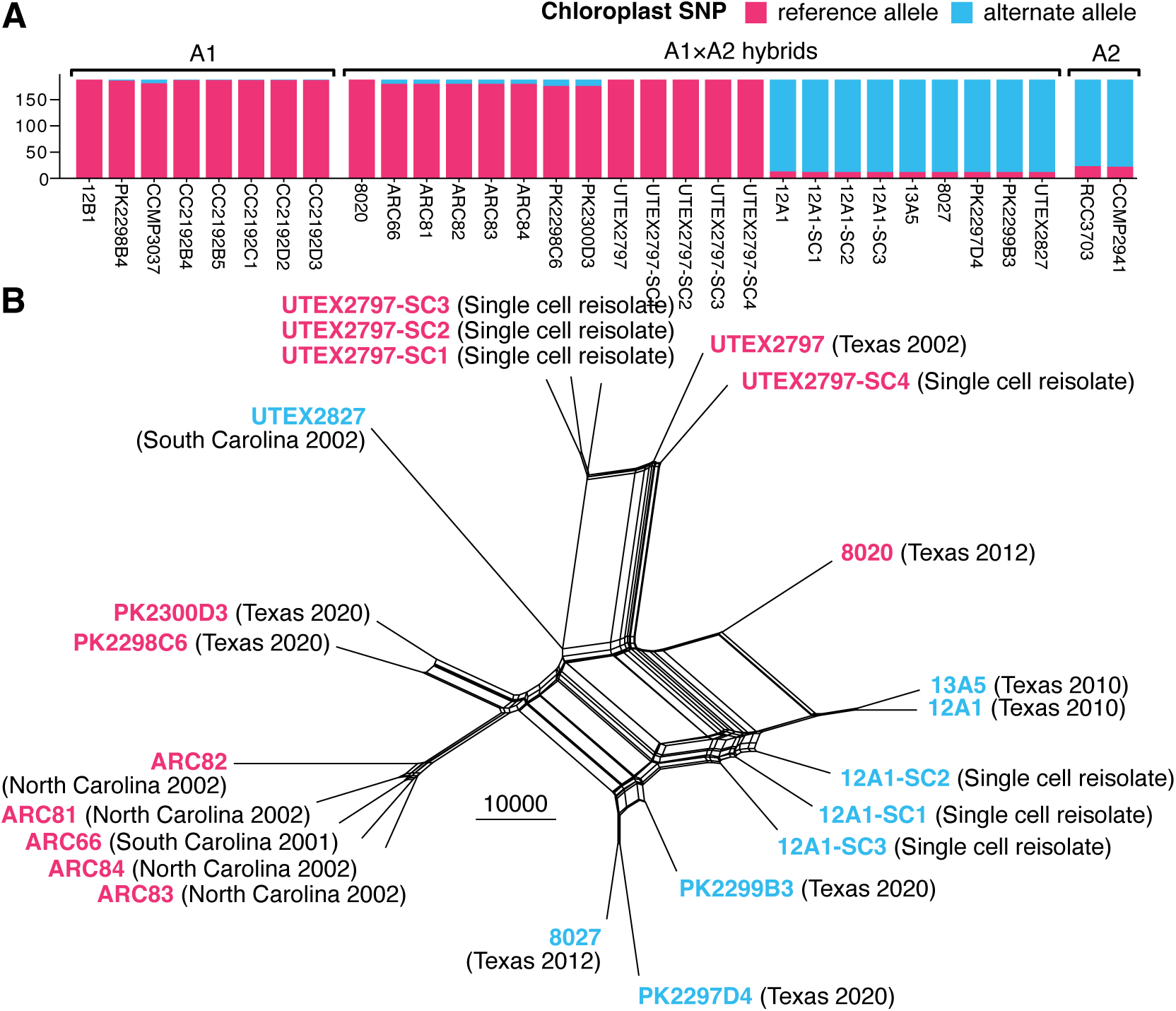
Genetic variation in A1×A2 hybrids. A) Chloroplast SNPs. B) Neighbor net analysis of A1×A2 hybrids: cpHap-A1 (pink), cpHap-A2 (blue). Edge lengths represent absolute SNP differences with the scale bar corresponding to 10,000 SNPs.

Strains with different cpHaps were co-isolated from the same blooms. In Lake Possum Kingdom, Texas, strains 8020 (cpHap-A1) and 8027 (cpHap-A2) were co-isolated from the same bloom in 2012. Hybrids with both cpHap-A1 (PK2298C6 and PK2300D3) and cpHap-A2 (PK2297D4 and PK2299B3) were also co-isolated in 2020. The three Lake Possum Kingdom strains belonging to cpHap-A2 cluster together in the nuclear SNP phylogenetic network, but the three cpHap-A1 strains do not (Figure 3B). Occurrence of cpHap-A1 and cpHap-A2 hybrid strains extends beyond Lake Possum Kingdom. Strain ARC66 (cpHap-A1) and UTEX2827 (cpHap-A2) were both isolated from water samples from South Carolina in 2001 and 2002, respectively.

The clear segregation of hybrid chloroplast variation is contrasted by a more complex pattern in the nuclear genome. An analysis of molecular variance (AMOVA) showed that cpHap explained only 22% of the variance present in the hybrid nuclear SNP dataset compared to 98% of the variance present in chloroplast SNPs (*p* < 0.001 for both). We also classified hybrid strains based on region of isolation, inland Texas (n=16) or coastal Carolinas (n=6), and found geography explained a similar amount of nuclear variance (26%, *p* < 0.001). The interaction between region and cpHap was explored further using hierarchical AMOVA (Table S4). When region was nested within cpHap, cpHap lost significance (Φ*_CT_* = -0.055, *p* = 0.344) while region within cpHap remained significant (Φ*_SC_* = 0.419, *p* < 0.001). The pattern was reversed when cpHap was nested within region (Φ*_CT_* = 0.069, *p* = 0.333 for region; Φ*_SC_* = 0.370, *p* < 0.001 for cpHap). The AMOVA results indicate that hybrid genome variation reflects both geography and chloroplast type, but that the two factors are partially overlapping rather than hierarchical.

### Chromosomal and segmental anomalies accumulate in laboratory hybrid strains

Chromosomal anomalies including aneuploidy (gain or loss of whole chromosomes) and large-scale structural mutations resulting in gain or loss of chromosomal segments have been reported in hybrid genomes of other microbial eukaryotes (van den Broek et al. 2015; Inbar et al. 2019; Maeda et al. 2022; Craig et al. 2025). To investigate whether similar chromosomal anomalies were present in A1×A2 hybrids, we plotted normalized coverage (NC; i.e., normalized read depth) alongside the relative abundance of A1- vs A2-derived reads. Numerous large-scale structural anomalies suggestive of duplications and losses of either A1 or A2 haplotypes were clearly visible in genome plots of hybrid strains (e.g., Figure 4). We chose to first compare strains UTEX2797 and 12A1 to their single-cell reisolates. This approach circumvents potential reference bias by treating each progenitor as its own baseline. Moreover, anomalies observed in reisolates have arisen *de novo* after the progenitor strains were first cultured in 2002 and 2012, respectively. We classified anomalies into three categories: chromosomal anomalies spanned whole scaffolds, telomeric anomalies involved the terminal ends of scaffolds, and interstitial anomalies were found in scaffold interiors.

**Figure 4:**
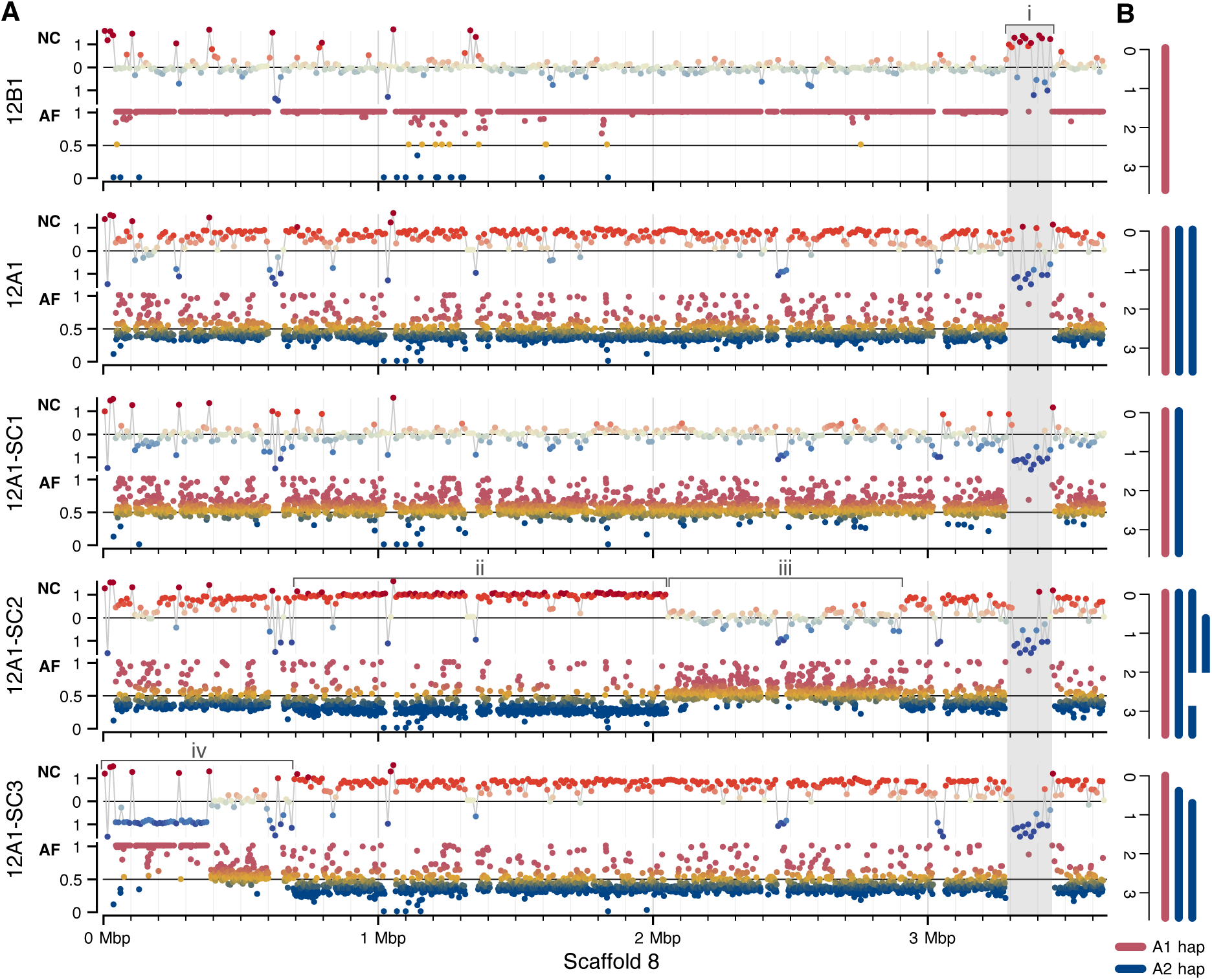
Scaffold 8 structural anomalies in 12A1 and single-cell reisolates. A) For each strain, top track indicates log_2_ ratio of normalized genome coverage (NC); bottom track indicates A1 AIM allele frequency (AF; A1 read depth / total read depth). Region i highlights a misassembly in the reference genome that was skipped in our analysis. Regions ii and iii correspond to interstitial anomalies in 12A1-SC2. Region iv indicates two overlapping telomeric anomalies in 12A1-SC3. B) Models of scaffold 8 organization indicating regions duplicated or lost in single-cell reisolates relative to 12A1.

In total, four chromosomal anomalies, four telomeric anomalies, and five interstitial anomalies were identified in single-cell reisolates (Table 2, Table S5, Figure S4-S9). Critically, genetic information absent in the progenitor culture remained absent in all single-cell reisolates. For example, Scaffold 22 in UTEX2797 has three large regions (two telomeric and one interstitial) with no heterozygosity; these regions showed a decrease in NC and loss of A2 alleles consistent with loss of the A2 haplotype. All UTEX2797 single-cell reisolates show the same loss; the A2 haplotype never reappears in these regions (Figure S8).

**Table 2:**
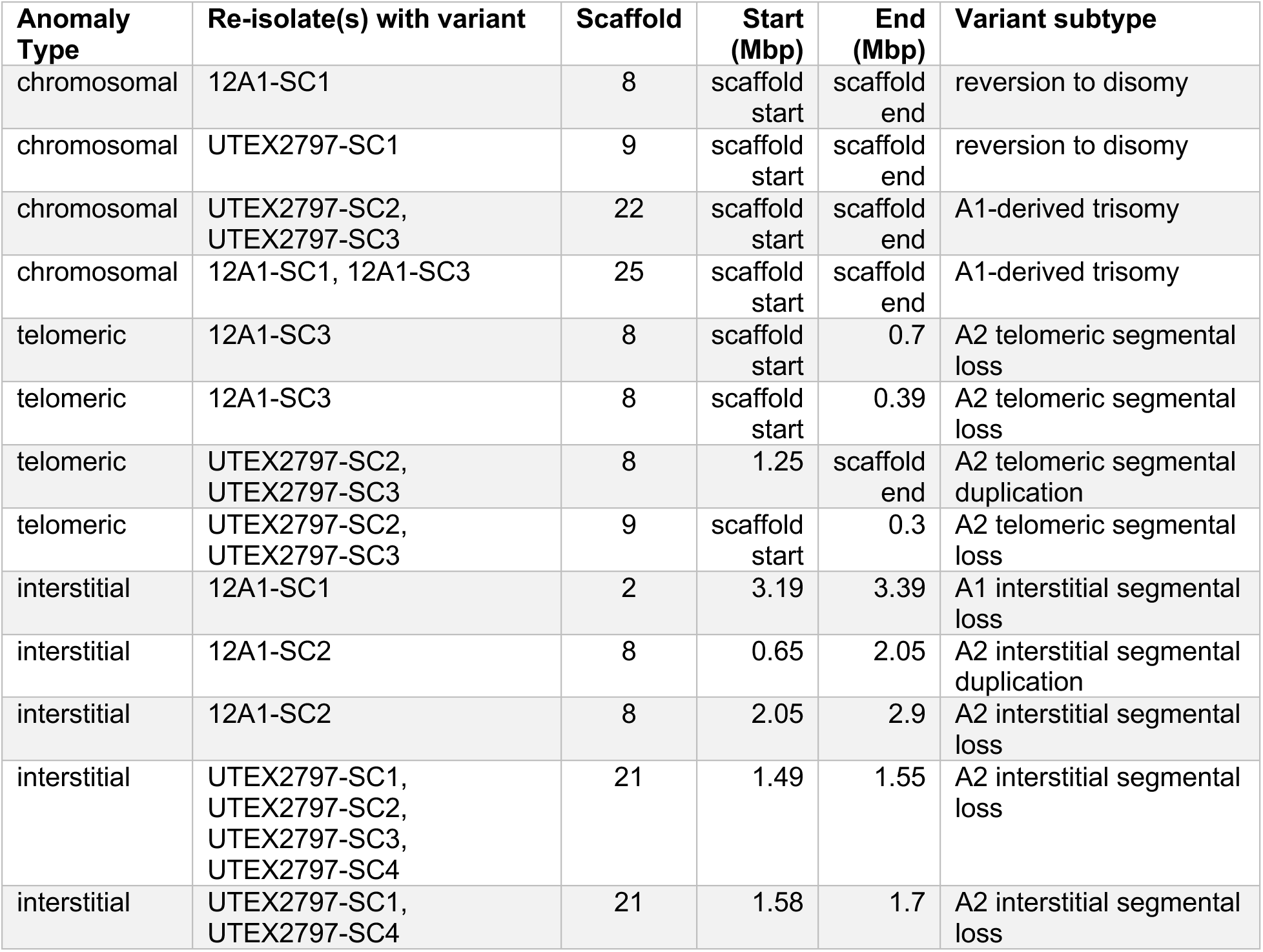
List of structural anomalies in single-cell reisolates compared to progenitor strains.

To illustrate the range and dynamics of anomalies observed, we describe Scaffold 8 in detail below, as it contains examples of all three categories across the progenitor strain, 12A1, and its reisolates (Figure 4). In 12A1, Scaffold 8 shows elevated A2 allele frequency (A2_AF) across its full length alongside an increase in NC, which is consistent with a duplication of the A2-derived chromosome. Single-cell reisolate 12A1-SC1 showed a decrease in both NC and A2_AF indicating a reversion to disomy. 12A1-SC2 contains two examples of interstitial anomalies: Scaffold8:650000-2050000 shows increased NC and A2_AF indicative of an A2 interstitial duplication; Scaffold8:2050000-2900000 shows decreased NC and A2_AF indicative of A2 interstitial loss (Figure 4). Lastly, 12A1-SC3 contains two examples of telomeric losses, both involving the beginning of its two A2-derived chromosomes (Figure 4).

### A1×A2 hybrids contain more genome anomalies than non-hybrid strains

We extended our analysis of genome structural anomalies to all *P. parvum* A1, A2, and A1×A2 hybrid strains, excluding single-cell reisolates, which were analyzed separately above. Across 24 strains, the median count of genome anomalies was 2.5 with seven strains (six non-hybrid and one hybrid) having no anomalies (Figure 5A). None of the diploid A1-type strains isolated in 2020 contained any anomalies. Anomaly counts were significantly higher in hybrid strains compared to non-hybrid strains (Wilcoxon-Mann-Whitney test, *Z* = 2.55, *p* = 0.0096; Figure 5B).

**Figure 5.**
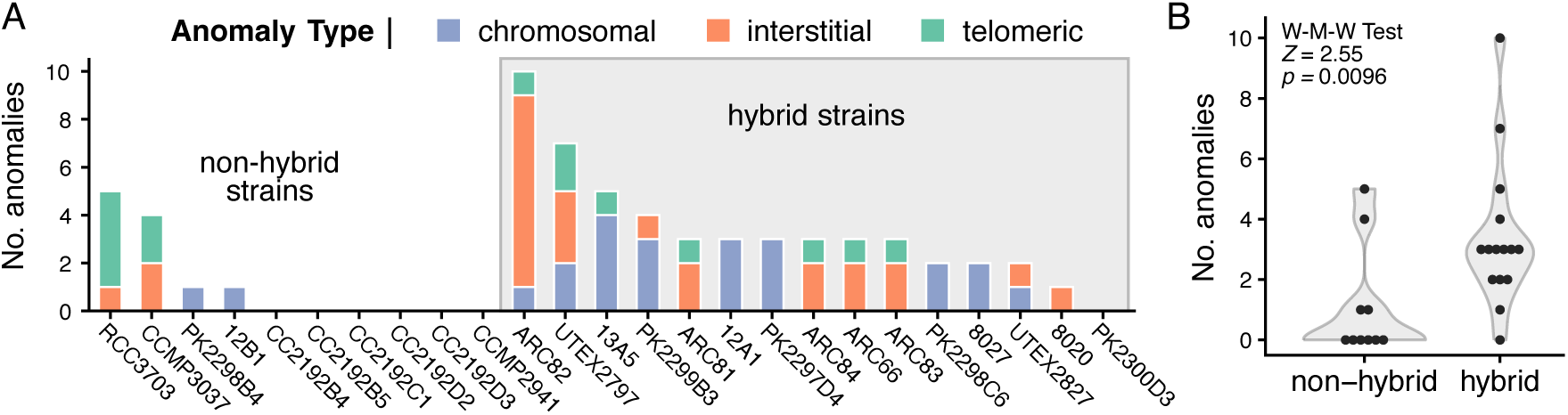
Structural anomalies detected in A1, A2, and A1×A2 strains. A) Number of chromosomal, interstitial, and telomeric anomalies detected per strain. B) Violin plots comparing hybrid vs non-hybrids. Statistical analysis was performed using a Wilcoxon-Mann-Whitney test.

The strain with the most anomalies was hybrid strain ARC82 (n = 10). Yet, this count represents an underestimate as Scaffold 28 in ARC82 displayed extreme variability in both NC and AF, which was indicative of significant chromosomal rearrangement relative to the 12B1 reference assembly, and prevented accurate inference of anomaly count for this scaffold in ARC82 (Figure S10). We struggled to accurately estimate the number of structural anomalies in a second strain, UTEX2827, which had two clear anomalies: a Scaffold 9 polysomy and an interstitial loss on Scaffold26 (Table S6). However, several other regions in UTEX2827 showed high variability NC and AF that prevented an accurate count of structural anomalies for this strain across several scaffolds (Figure S11).

Unlike anomalies in single-cell reisolates, whose timing of origin is constrained by the known isolation dates of the progenitor strains, the evolutionary timing of anomalies detected in the remaining strains is often uncertain. Some anomalies may have evolved in lab cultures, whereas others may represent genomic variation present in natural populations. Four cpHap-A1 strains from North and South Carolina (ARC66, ARC81, ARC83, and ARC84) share the same three structural anomalies: an A2 interstitial loss on Scaffold 4, an A1 telomeric loss on Scaffold 13, and an A1 interstitial duplication on Scaffold 26 (Table S6). Similarly, all five cpHap-A2 strains from Texas (8027, 12A1, 13A5, PK2297D4, and PK2299B3) are A1-disomic for Scaffold 27, having lost the A2-derived copy (Table S6). The presence of the same structural anomalies in closely related but independently isolated strains suggests these genetic variants were present in the natural population from which they were sampled. In contrast, ARC82 was sampled from the same location and in the same year as ARC81, ARC83, and ARC84 but shares only one anomaly (the A1 interstitial duplication on Scaffold 26) with these strains (Table S6). Other closely related strains (e.g., PK2298C6 and PK2300D3; 12A1 and 13A5; and PK2297D4 and PK2299B3) also differ in some anomalous regions (Table S6). Whether these differences represent standing variation present in natural populations or culturing-induced differences requires further study.

Lastly, two scaffolds show evidence for repeated polysomies (Table S6). Scaffold 8 was polysomic in 20.8% of strains in our analysis: four cpHap-A2 strains (12A1, 13A5, PK2297D4, and PK2299B3) and one cpHap-A1 strain (PK2298C6). This polysomy is notably absent in PK2300D3 (closely related to PK2298C6) and 8027 (closely related to the four cpHap-A2 strains), suggesting that polysomy of Scaffold 8 is either recurrent or frequently lost. This is consistent with the pattern observed in 12A1-SC1, which reverted to disomy from the polysomic state present in 12A1 (Figure 4). Similarly, Scaffold 19 was polysomic in 37.5% of strains in our analysis: five cpHap-A2 strains (8027, 12A1, 13A5, PK2297D4, PK2299B3), two cpHap-A1 strains (PK2298C6, ARC82), and two A1 strains (PK2298B4, 12B1). The presence of Scaffold 19 polysomies in both hybrid and non-hybrid strains suggests that some chromosomes may be prone to polysomy regardless of hybrid status, though whether this reflects a shared ancestral variant or recurrent independent events remains unclear.

## DISCUSSION

Strains of *P. parvum* s.l. have been studied in laboratory cultures for over 70 years and have never been observed to reproduce sexually. Nevertheless, the existence of A1×A2 hybrid strains provides strong evidence for a facultative sexual lifecycle in these algae. The lifecycles of other haptophyte algae are poorly understood outside of the coccolithophores, which have complex sexual haplodiplontic life cycles (Bousquet et al. 2025). In coccolithophores, lifecycle phases are heteromorphic; cells alternate between a flagellated haploid phase and a calcified diploid phase, each capable of clonal reproduction. *P. parvum* s.l. likely undergoes a similar haplodiplontic lifecycle, but the haploid and diploid phases are morphologically indistinguishable (Figure 6). Different A1×A2 hybrid strains carry chloroplast haplotypes from either the A1 or A2 parent, indicating that while chloroplast inheritance is uniparental during syngamy, both parental lineages can serve as the chloroplast donor (Figure 6). Additional work is needed to determine whether distinct mating types exist in *P. parvum* s.l. as in other algae, such as *Chlamydomonas* (Umen and Coelho 2019). In the past, research on the *P. parvum* s.l. lifecycle has been complicated by the cryptic nature of the species complex, both in terms of cryptic divergence between clades and cryptic lifecycle phases. Identification of multiple haploid and diploid strains within the A1 clade (Table 1) will facilitate new research into the *P. parvum* s.l. lifecycle.

**Figure 6:**
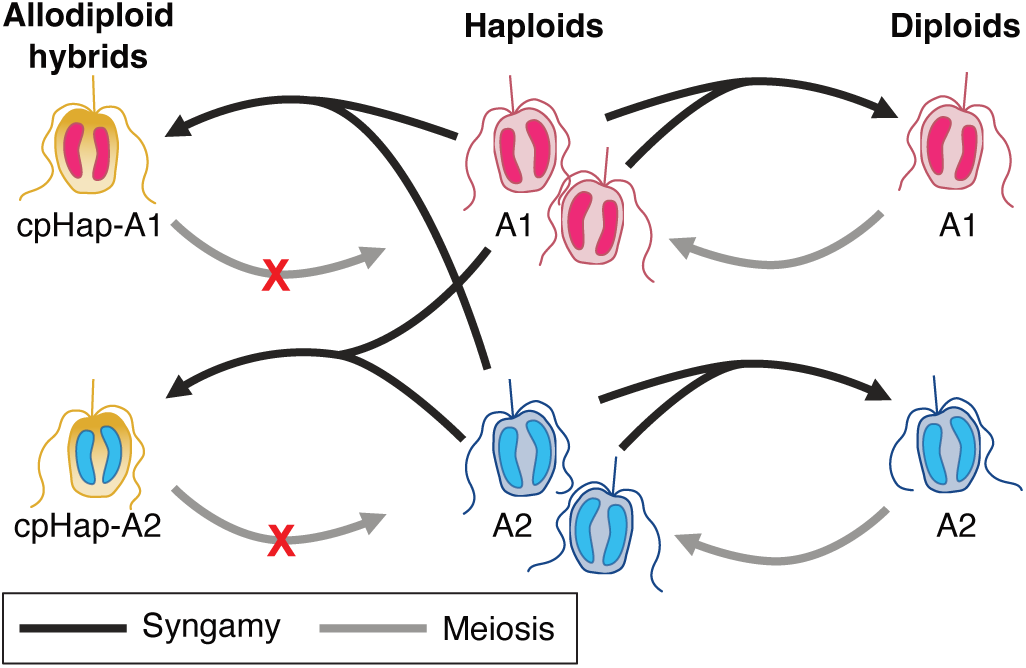
Proposed model of *P. parvum* clade A lifecycle and hybridization. Allodiploid hybrids are formed via syngamy between A1 and A2 haploid cells; genetic incompatibilities prevent meiosis (red X). During hybridization, chloroplasts can be inherited from either the A1 parent (cpHap-A1, pink) or the A2 parent (cpHap-A2, blue). All cell types are capable of asexual reproduction via mitosis (not shown).

The lack of intermediate strains that group in between the hybrid clade and either the A1 or A2 clades (Figure 2A) suggests a lack of backcrossing between hybrids and their parent populations. One possible explanation for this pattern could be that A1×A2 hybrids cannot complete meiosis (Figure 6) as is the case in some other hybrid systems (Bizzarri et al. 2016; Montecinos et al. 2017). This could be due to divergent parental karyotypes such that homologous chromosomes are unable to pair up correctly during meiosis (Hunter et al. 1996). Alternatively, genetic incompatibilities between the A1 and A2 alleles could lead to low viability of cells post meiosis (Xu and He 2011). The large number of chromosomal abnormalities present in our existing hybrid lab strains would certainly reduce the number of viable gametes produced following meiosis, but the exact timing of when most abnormalities evolved is unclear. In our phylogenomic analysis of protein-coding genes, the A1 and A2 clades clearly segregate with strong branch support (Figure 2B), consistent with the accumulation of significant genetic differences since their divergence. Additional chromosome-scale genome assemblies and comparative karyotyping of *P. parvum* clade A are needed to investigate whether the A1 and A2 strains possess chromosomal number differences and/or structural rearrangements that would inhibit meiosis in hybrids. While A1×A2 hybrids appear reproductively isolated from their parental populations, examples from yeasts suggest that hybrid fertility can sometimes be evolutionarily rescued through mutation of mating type loci (Ortiz-Merino et al. 2017; Braun-Galleani et al. 2018), raising questions about the long-term evolutionary fate of *P. parvum* clade A hybrids.

The hybrid strains in our analysis are likely the products of multiple A1×A2 hybridization events. The existence of two chloroplast haplotypes is a strong indication that hybridization has occurred at least twice. However, if hybrid strains were the result of only two events (one event giving rise to cpHap-A1 strains and the other event producing cpHap-A2), that should be reflected in the nuclear data as well. Nuclear variation does cluster based on cpHap, but only after accounting for geography (Table 2). This pattern is more consistent with independent hybridization events producing cpHap-A1 and cpHap-A2 strains in Texas separate from those that gave rise to the same cpHap in the Carolinas.

Despite most strains in our analysis being from North America, *P. parvum* type A has a broad global distribution with cataloged strains from Europe, Israel, Japan, and Australia (Binzer et al. 2019). In our analysis, Strain NIES1812 from Okinawa, Japan grouped separately from the A1 and A2 clades, consistent with earlier work (Binzer et al. 2019), indicating that the strain is sufficiently divergent to warrant designation as a separate A3 subclade. Overall, the *P. parvum* type A subclades show some geographic clustering, with A1 strains from North America, A2 strains from Europe, and the single A3 clade from Japan. However, this pattern should be interpreted cautiously, as *P. parvum* type A distribution and diversity remain poorly understood even in North America, where sampling has been most extensive.

The possibility of multiple hybridization events in *P. parvum* clade A raises the intriguing prospect of one or more hybrid zones where A1 and A2 populations overlap. Hybrid zones are well documented in plants and animals (Harrison 1993), but less common in microbial eukaryotes. This may be due to difficulty in detection rather than biological barriers to hybridization. Here, *P. parvum* A1×A2 hybrids are morphologically indistinguishable from nonhybrid strains and must be identified based on genetic characteristics. Moreover, A1- and A2-type strains are also morphologically identical to one another, and the true range of either parental population is unknown. Despite limited sampling, we know that different lineages of *P. parvum* type A can co-occur in nature. A1 and A1×A2 hybrid strains were co-isolated from the same water sample twice in Texas: once in 2010 from a bloom in Moss Creek Lake (Driscoll et al. 2013) and again in 2020 from a bloom in Possum Kingdom Lake (Table 1). Characterizing the genetic diversity of *P. parvum* s.l. present at a location based on a limited number of isolated strains may give a biased representation of true diversity. Culture-independent methods, such as single-cell sequencing, could be employed in future studies of *P. parvum* s.l. to determine where or if hybrid zones exist.

Rapid evolution of extensive segmental and chromosome-scale anomalies appears to be a common feature of hybrid genomes in microbial eukaryotes, with precedents in fungi, diatoms, trypanosomatids, and green algae (van den Broek et al. 2015; Inbar et al. 2019; Maeda et al. 2022; Craig et al. 2025). These anomalies can have functional consequences. For example, in lager brewing yeast strains, different aneuploid variants among hybrids were associated with distinct phenotypes in flocculation and diacetyl production (van den Broek et al. 2015). Heterosis, also known as hybrid vigor, where hybrid strains show increased growth compared to parental strains has been documented in natural hybrid fungal endophytes (Li et al. 2024) and demonstrated experimentally in *Saccharomyces* spp. (Bernardes et al. 2017). Conversely, some variants may be maladaptive or only tolerated under controlled laboratory conditions. Similar to observations of the green algal *Auxenochlorella* hybrid (Craig et al. 2025), we observed high levels of structural anomalies between single-cell isolates and their progenitor cultures, suggesting that genome instability may accumulate more readily in stable culture conditions than in natural environments. However, our analysis also identified anomalies shared between independently isolated strains, indicating some aneuploid and segmental variation is present in natural populations of *P. parvum.* Future studies are needed to determine how many of the chromosomal abnormalities observed in lab strains are also present in natural populations.

The number of structural anomalies identified in *P. parvum* clade A strains was striking and indicates that substantial genomic diversity exists within this group. Moreover, our estimates represent a lower bound on the true extent of structural variation present in laboratory strains. Reference bias is one important limitation as any sequence absent in 12B1, including A2-specific regions, will go undetected in our analysis. Additionally, we could only characterize anomalies as losses or duplications relative to the reference genome and could not characterize other types of structural variation such as translocations or inversions. Our results underscore the need for additional chromosome-scale genome assemblies, including from A2 strains and A1xA2 hybrids, to fully reconstruct genome structural differences present in *P. parvum* clade A.

The functional and ecological consequences of hybridization in *P. parvum* s.l. remain poorly understood, yet may be directly relevant to predicting and managing *P. parvum* s.l. harmful algal blooms given the prevalence of hybrid strains in North American bloom samples. Phenotypic variation is known to exist between clade A strains, including differences in prymnesin profiles (Binzer et al. 2019) as well as growth rates and feeding behavior among A1 and hybrid strains (Driscoll et al. 2013; Driscoll et al. 2023). No A-type prymnesin producing strain has been more extensively studied than hybrid strain UTEX2797, which has served as a model strain for the clade in numerous experiments. Notably, A-type prymnesins from UTEX2797 were found to be the most potent among tested strains (Varga et al. 2024), but two extracts from UTEX2797 sampled three years apart showed significant differences in prymnesin profiles and potency. Our findings suggest that genome instability during laboratory culture could contribute to such variation, raising important questions about the reproducibility of studies conducted across time with the same strain. More broadly, it remains unclear how representative observations made with UTEX2797 are of non-hybrid strains, or whether hybrid and non-hybrid populations differ in their propensity to form toxic blooms. Given the ongoing threat of *P. parvum* s.l. blooms to freshwater systems worldwide (Roelke et al. 2016; Macêdo et al. 2023) and the substantial genomic variation documented here among laboratory strains, understanding the relationship between genetic diversity, hybridization, and toxicity is an important topic for future investigation.

## MATERIALS AND METHODS

### Taxon sampling

Illumina sequence data was acquired for 38 *P. parvum* strains from cultures in the Wisecaver Lab (see DNA sequencing, read filtering, and read mapping). Of these strains, 28 were from culture collections, including the Algal Resource Collection (6 strains courtesy of Dr. Schonna Manning); the National Center for Marine Algae and Microbiota (2 strains); the Culture Collection of Algae at the University of Texas at Austin (5 strains, 2 courtesy of Dr. William Driscoll); the Norwegian Culture Collection of algae (3 strains courtesy of Dr. Tim Fallon); the Roscoff Culture Collection (5 strains); the Kalmar Culture Collection (1 strain courtesy of Dr. Tim Fallon), and the Microbial Culture Collection at the National Institute of Environmental Studies, Japan (3 strains). Three additional strains were acquired from the personal collection of Dr. William Driscoll. See Supplemental Table S1 for additional strain information.

The remaining 10 strains were isolated from 2019 and 2020 *P. parvum* s.l. blooms in Texas. Water samples from a 2019 bloom in Lake Colorado City and a 2020 bloom in Possum Kingdom Lake were collected by the Texas Parks and Wildlife Department (TPWD) as part of the agency’s Golden Alga monitoring program and mailed courtesy of Greg Southard. To generate clonal cultures, water samples were passed through a MoFlo Astrios High Speed Cell Sorter (Beckman Coulter, Brea, CA, USA). A gated population was identified based on forward scatter and chlorophyll fluorescence of existing *P. parvum* s.l. pure culture, and positive events were sorted into 96-well plates, one event per well. To create the initial plate media, TPWD water samples were passed through a 0.22 μm polyethersulfone filter (CELLTREAT, Pepperell, MA, USA). Plates were grown at 20°C using a 12:12 light:dark cycle and irradiance of approximately 100 µmol photos m^-2^ s^-1^. After three weeks, any well consisting of *P. parvum* was retained as an isolate and transferred to 20 mL of L1-Si media in sterile 25 x 150 mm borosilicate glass test tubes. This resulted in 5 new isolates from the Lake Colorado City bloom and 5 new isolates from the Possum Kingdom Lake bloom.

### Algal culturing

L1-Si media was prepared using Instant Ocean Sea Salt (Spectrum Brands, Blacksburg, VA, USA) at a salinity of 6 and a L1-Si Media Kit obtained from the National Center for Marine Algae and Microbiota (East Boothbay, ME, USA). All cultures were maintained in a 12:12 light:dark cycle at 20°C. Media and salinities used for each strain can be found in Table S1.

Cultures were periodically treated with antibiotics to reduce bacterial contamination (Guillard 2005). Two milliliters of dense (>5x10^5^ cells/mL) late-exponential phase cultures were transferred sequentially into media containing different antibiotics: first ampicillin (100 mg/L) and neomycin (100 mg/L) for 48 hours, followed by kanamycin (100 mg/L) and penicillin (100 mg/L) for another 48 hours, and chloramphenicol (100 mg/L) for a final 24 hours. Subsequently, 2 mL of culture was transferred to fresh L1-Si media and allowed to fully recover.

### Single-cell reisolation

UTEX2797 and 12A1 cultures were grown to late exponential phase before single-cell reisolation via serial dilution, following established protocols for single-cell isolation of microalgae (Andersen and Kawachi 2005; Lu et al. 2025). Cells were serially diluted to 1 cell/mL in L1-Si media (salinity 6) and transferred to 96-well plates with 100 μL of diluted culture per well. Cultures were allowed to recover, and wells were considered derived from a single cell if ≤ 9 wells per plate had growth.

### DNA sequencing, read filtering, and read mapping

Cultures for DNA sequencing were grown in 100 mL L1-Si media in 250 mL glass Erlenmeyer flasks. For each sample, 50 mL of late-exponential phase *P. parvum* s.l. culture was spun down at 4000 x g for 10 minutes. Genomic DNA was extracted from cell pellets using Plant DNAzol (Thermo Fisher #10978021, Waltham, MA, USA) according to lab protocol (Wisecaver 2026). Sequencing libraries were constructed and sequenced to produce 2x150 bp paired-end reads using an Illumina NovaSeq 6000 by Novogene Corporation Inc. (Sacramento, CA, USA).

Illumina read quality was assessed with FastQC v0.11.9 (Babraham Bioinformatics 2011) and reads were trimmed as needed using FastP v0.20.1 with quality, low complexity, polyG, and polyX filtering enabled (Chen et al. 2018). Although antibiotic treatments were performed as necessary, all cultures were considered xenic. Therefore, reads derived from contamination were identified and removed via the following pipeline. For each sample, reads were first assembled with ABySS v2.3.1 (Simpson et al. 2009) using a k-mer size of 96. Assembly scaffolds were queried against the NCBI nucleotide (nt) database (accessed September 11^th^, 2021) using blastn v2.11.0+ (Camacho et al. 2009) and against a custom RefSeq database (O’Leary et al. 2016) using DIAMOND v2.0.8 (Buchfink et al. 2015). The custom RefSeq database was composed of RefSeq release 207 supplemented with predicted protein sequences from marine microbial eukaryotic transcriptome sequencing project (MMETSP) (Keeling et al. 2014) and the 1000 plants transcriptome sequencing project (1KP) (Matasci et al. 2014). Reads were aligned against the ABySS assembly using minimap2 v2.24-r1122 (Li 2018). Assembly scaffolds were filtered using BlobTools v1.1.1 (Laetsch and Blaxter 2017). The BlobTools taxrule ‘bestsumorder’ determined the taxonomic assignment of each scaffold, prioritizing information from protein hits first. Scaffolds denoted as non-eukaryotic in origin were removed to produce a filtered assembly. BBSplit v38.87 from the BBTools software suite (Bushnell 2017) was used to retain reads which mapped to the filtered assembly (hereafter referred to as filtered reads).

Filtered short reads for all strains were aligned to the 12B1 v1.1 genome (GenBank assembly GCA_041296205.1; Wisecaver et al. 2023; Fallon et al. 2024) using BWA-MEM v0.7.17 (Li 2013). Read alignments were filtered to include only reads mapped in proper pairs with a minimum quality of 30 via SAMtools v1.9 (Li et al. 2009; Danecek et al. 2021). Secondary and chimeric alignments flagged by BWA-MEM v0.7.17 (Li 2013) were also removed. PCR duplicates were removed with GATK MarkDuplicates v4.2.6.1 (McKenna et al. 2010). Median read alignment rate was 90.4% and ranged from 78.2% (ARC82) to 96.6% (CCMP3037) (Table S7).

### Preliminary SNP phylogenetic network

A preliminary SNP network was constructed to first identify all *P. parvum* clade A strains for downstream analyses (see analysis flowchart Figure S8). Variants were called against the 12B1 v1.1 genome using BCFtools v1.17 mpileup (Danecek et al. 2021) using BAM alignment files for all 38 strains (excluding single-cell reisolates) as input (Table S1). Variants were first quality filtered using VCFtools 0.1.16 (Danecek et al. 2011) to retain only biallelic SNPs with a minimum quality of 100 and a minimum p-value of 1x10^-6^. For constructing the phylogenetic network, biallelic SNPs were further filtered using VCFtools so that the final dataset included those present in all individuals, a minimum mean depth of 20, and a minimum minor allele frequency of 0.04 (i.e. at least two samples). The resulting 816,587 SNPs were used to generate a square distance matrix with PLINK v1.90b6.21 (Purcell et al. 2007). A network depicting this matrix was created using NeighborNet with default options in SplitsTree4 v4.19.2 (Huson 1998; Bryant 2003). Strains that grouped with strains known to produce A-type prymnesin (Binzer et al. 2019; Wisecaver et al. 2023) were included in all downstream analyses, with the exception of ARC140. ARC140 was originally chemotyped as producing B-type prymnesin (Binzer et al. 2019), but clustered with type A strains in our analysis, so it was excluded.

### Genome size estimation via flow cytometry

Exponentially growing *P. parvum* s.l. culture (500 µL) was centrifuged at 1000 x g for 10 minutes. The supernatant was discarded and the pellet flash-frozen in liquid nitrogen and resuspended in 500 µL ice-cold nuclear isolation buffer LB01 consisting of 15 mM tris, 2 mM Na_2_EDTA, 0.5 mM spermine tetrahydrochloride, 80 mM KCl, 20 mM NaCl, and 0.1% (v/v) Triton X-100 at pH 8.0 (Dpooležel et al. 1989; Čertnerová and Galbraith 2021). Nuclei were kept at 4°C and analyzed within 24 hours.

All nuclei were treated with 0.4 mg/mL RNAseA (Invitrogen #12091021, Waltham, MA, USA) and 0.05 mg/mL propidium iodide (Invitrogen #P3566, Waltham, MA, USA) at 4°C for 3 hours in the dark. Following incubation, nuclei were passed through a 35 µm cell-strainer cap (Falcon #352235, Corning, NY, USA) and analyzed on a Cytek Northern Lights flow cytometer equipped with a 488-nm laser and 14 fluorescent detection channels (Cytek Biosciences, Fremont, CA, USA). Nuclei were identified via forward scatter and fluorescence in the B7 channel corresponding to peak propidium iodide fluorescence. Fluorescence values corresponding to at least 1000 nuclei per sample were gathered using the batch analysis feature in SpectroFlo (Cytek Biosciences, Fremont, CA, USA). Biological replicates were performed on separate days at approximately the same time. DNA content of 12B1, RCC3703, CCMP3037, UTEX2797, 12A1, and K0081 from Wisecaver et al., (2023) were used to create a standard curve to convert fluorescence into total DNA content in picograms of DNA for all other strains (Table S2). Total DNA content was converted to base pairs using the formula of 1 pg = 0.98 x 10^9^ base pairs (Cavaller-Smith 1985).

### Estimation of heterozygosity

K-mers were counted from filtered reads using KMC v3.1.1 with a k-mer length (-k) of 21, minimal k-mer occurrence (-ci) of 1, and maximal k-mer occurrence (-cs) of 10,000 (Kokot et al. 2017). K-mer frequencies were modeled using GenomeScope2 v2.0 (Ranallo-Benavidez et al. 2020) using a k-mer length of 21 and a ploidy value of 1,2, or 4 depending on the strain (see Table 1). Strains with heterozygosity above 1% and a pronounced heterozygous peak were flagged as high heterozygosity.

### *P. parvum* clade A phylogenetic analyses

Variants were called against the 12B1 v1.1 genome using BCFtools v1.17 mpileup (Danecek et al. 2021), as described above using BAM alignment files for all clade A strains and single-cell reisolates (Table S1). Variants were filtered using VCFtools 0.1.16 (Danecek et al. 2011) and used to create a SNP phylogenetic network as described above (see analysis flowchart Figure S9). The resulting A-type only dataset was comprised on 453,986 SNPs.

To construct a gene-based species tree, cleaned short-read DNA libraries from non-hybrid A-types, B-types, and C-types were assembled into contigs via ABySS v2.3.1 (Simpson et al. 2009). Assemblies were masked with RepeatMasker v4.1.1 (Smit et al. 2017) using the modeled repetitive element library generated by Wisecaver et al., (2023). This masked genome was then used to create a hints file with BRAKER v2.1.5 (Hoff et al. 2019; Brůna et al. 2021) supplemented with evidence from Odb10-protozoa and all predicted haptophyte proteins from MMETSP (Keeling et al. 2014; Kriventseva et al. 2019). This hints file and an AUGUSTUS species-specific training file from Wisecaver et al., (2023) was used to predict coding sequences on all masked genomes with BRAKER v2.1.5 (Hoff et al. 2019; Brůna et al. 2021). The longest coding sequence for each locus was extracted using a custom python script and used input for OrthoFinder v2.5.5 (Emms and Kelly 2015; Emms and Kelly 2019). OrthoFinder identified 9836 single copy orthogroups present in all strains. Nucleotide coding sequences were aligned with MAFFT in the GUIDANCE v2.02 software suite (Katoh and Standley 2013; Sela et al. 2015). Alignments were trimmed using trimAl v1.4.1 (Capella-Gutierrez et al. 2009). All alignments smaller than 150 bp were discarded, leaving 9769 orthogroups for species tree construction. All alignments were concatenated and used to create a concatenated species tree via IQTree2 v2.2.2.9 (Nguyen et al. 2015; Minh et al. 2020). Size-filtered alignments were also used to create individual gene trees using IQTree2 v2.2.2.9 (Nguyen et al. 2015; Minh et al. 2020). The coalescence-based phylogeny estimation was conducted using ASTRAL v5.7.1 (Zhang et al. 2018). Gene and site concordance values were calculated for both the coalescent and concatenated species trees via IQTree2 v2.2.2.9 and visualized via FigTree v1.4.4 (Nguyen et al. 2015; Rambaut 2018; Minh et al. 2020). See Figure S14 for analysis flowchart.

### Identification of Ancestry Informative Markers

Ancestry Informative Markers (AIMs) diagnostic of A1 or A2 ancestry were identified as previously described (Blom et al. 2024; Thörn et al. 2024). Sites that were fixed (i.e., F_ST_ = 1) between diploid A1 genomes (parental population 1) and A2 genomes (parental population 2) were identified with the Weir and Cockerham method using VCFtools v0.1.16 (Danecek et al. 2011). This identified 277,926 AIMs. The F_ST_ was then calculated between each parental population and all other clade A strains individually, and AIMs in these strains were classified based on the difference between the two F_ST_ values. An AIM was classified as A1/A1 homozygous if F_ST__vs_A1 = 0 and F_ST__vs_A2 = 1. An AIM was classified as A2/A2 homozygous if F_ST__vs_A1 = 1 and F_ST__vs_A2 = 0. Lastly, an AIM was classified as A1/A2 heterozygous if both values were non-zero and the absolute value (abv) of the difference was less than 0.2; no AIMs had intermediate values (abv > 0.2 and < 1).

### Chloroplast SNP analyses

To identify reads that mapped preferentially to chloroplast genome, the 12B1_v1.1 nuclear genome was combined with the 12B1 chloroplast genome assembly from Wisecaver et al. (2023). Filtered short reads from A1, A2, and A1×A2 strains were aligned to the combined assembly using BWA-MEM v0.7.17 (Li 2013). Alignments were filtered to only include those mapped in proper pairs with a minimum quality of 30 via SAMtools (Li et al. 2009; Danecek et al. 2021). Supplementary and chimeric alignments as marked by BWA-MEM v0.7.17 (Li 2013) were also removed. PCR duplicates were removed with GATK MarkDuplicates v4.2.6.1. Variants were called using BCFtools v1.17 (Danecek et al. 2021) with minimum mapping and base qualities of 20. Variants were filtered using VCFtools 0.1.16 (Danecek et al. 2011) to only retain biallelic SNPs present in all individuals with a minimum quality of 30, a minimum minor allele frequency of 0.05, and a minimum mean depth of 20. See Figure S15 for analysis flowchart. The resulting chloroplast dataset consisted of 188 SNPs

To assess the partitioning of nuclear genetic variation among chloroplast haplotypes and geographic region of isolation, we conducted AMOVAs on the nuclear 453,986 SNP dataset and the chloroplast SNP dataset. AMOVAs were performed using the pegas function implemented in the poppr package (Kamvar et al., 2014) in R. Statistical significance of variance components was assessed by permutation tests (n = 999).

### Structural variant identification

Normalized genome coverage (NC) was calculated for each A1, A2, and A1×A2 strain with deepTools v3.5.5 (Ramírez et al. 2016) using the filtered BAM alignment files as input, bin size of 10 kbp, RPGC normalization, and an effective genome size of 93,538,723 (corresponding to the length of the 12B1_v1.1 assembly). Uniformity of genome coverage was evaluated, and the five smallest scaffolds 30-34 were excluded from the coverage analysis due to increased variability in coverage both within and between strains. A1 diploid strains CC2192B5 and CC2192D3 were selected to as reference euploid strains, as these strains had the lowest coefficient of variation in coverage across chromosomes and no outlier chromosomes after scaffolds 30-34 were excluded. The log_2_ ratio of NC for each 10 kbp window was calculated as log_2_NC = log_2_(x_i_ + 0.01) − log_2_(μ_diploid_ + 0.01), were x_i_ was the NC for strain *i* in a bin and μ_diploid_ was the mean NC for the reference strains in the same bin.

A1 allele frequency (A1_AF) was calculated for all AIMs by extracting the A1 allele depth from the VCF file divided by the total read depth for that site. The mean log2NC for 50 kbp windows and mean A1_AF for 20 kbp windows were plotted using Matplotlib (Hunter 2007). Diploid regions that have maintained both A1 and A2 parental alleles are expected to have an average log_2_NC ≈ 0 and A1_AF ≈ 0.5. Large regions that deviate from these expected values are indicative of structural variation. For example, a monosomic region resulting from a loss of the A1 haplotype would have half the coverage of diploid regions (log_2_NC ≈ -1) and a corresponding drop in A1_AF to zero; a trisomic region resulting from a duplication of the A1 haplotype would have 1.5x coverage (log_2_NC ≈ 0.58) and a corresponding 2-to-1 A1-to-A2 allele ratio (A1_AF ≈ 0.67). However, these numbers represent ideal values. Actual strain data displayed significantly more noise due to variation in read mapping quality across the genome and between haplotypes. Due to genome reference bias, reads derived from the A1 haplotype aligned better than A2 haplotype-derived reads. This resulted in elevated average A1_AF (≈ 0.57) in background regions of the genome (Table S3). Plots were manually inspected to identify regions of the genome (∼50 kbp or larger) that deviated significantly from log_2_NC = 0 and A1_AF = 0.57.

### Disclosure of AI use

The AI tool claude.ai versions Sonnet 4.5 and 4.6 were used to generate code for BAM and VCF processing, data analysis, and figure generation (Figures 3B, Figure 4, Figure S4-S6, and Figure S8). All code is available through GitHub (see Data Availability) and was reviewed, tested, and validated by the authors to ensure accuracy. The authors take full responsibility for the accuracy of all figures generated with the assistance of AI.

## DATA AVAILABILITY

Raw sequencing reads have been deposited in the Sequence Read Archive database under accession number PRJNA807128. VCF files, network files, tree and alignment files, and other related data files are available through FigShare (10.6084/m9.figshare.32572215). Scripts are available through GitHub (https://github.com/WisecaverLab/Pparvum_hybrid_diversity/). Strains are available upon request.

## ACKNOWLEDGEMENTS

We thank members of the Wisecaver lab and Dr. Brian Dilkes for helpful discussions. We thank Greg Southard of Texas Parks and Wildlife for sending water samples of *P. parvum* s.l. blooms. We thank Dr. Schonna Manning, Dr. Tim Fallon, and Dr. William Driscoll for sharing of several strains analyzed in this study. We also thank Raeya Ogas, Grace Estep, Olivia Riedling, and Dr. Bob Auber for their assistance with DNA isolation. This work was conducted in part using the resources of the Center for Institutional Research Computing at Washington State University. This work was also conducted in part using the resources of the Rosen Center for Advanced Computing at Purdue University and well as through use of the Flow Cytometry and Cell Separation Facility at the Bindley Bioscience Center at Purdue, a core facility of the NIH-funded Indiana Clinical and Translational Sciences Institute. This work was supported by the National Science Foundation under grant DEB-1831493 to JHW.

