## Supplemental Figures for "Hybridization between toxic bloom-forming algae of the *Prymnesium parvum sensu lato* species complex"

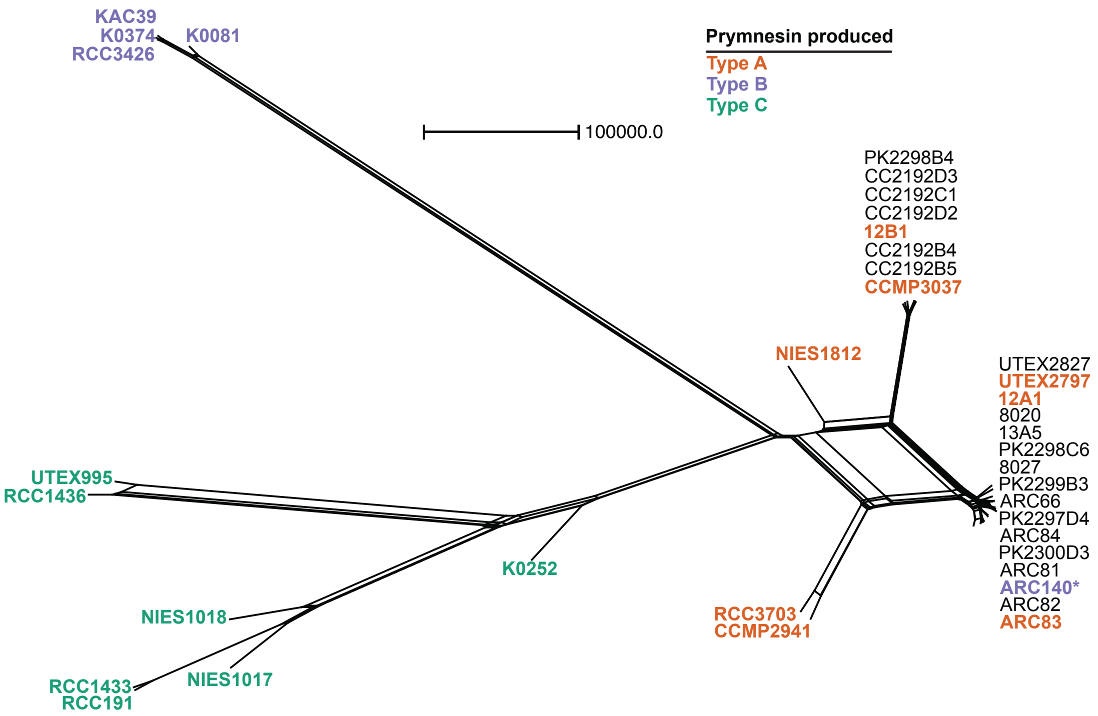


**Figure S1.** ***Prymnesium parvum*** **s.l. SNP network.** Chemotyped strains are color coded according to Wisecaver et al. (2023) and Binzer et al. (2019). Strains that grouped within the type A clade were included in all downstream analyses, with the exception of ARC140(*). ARC140 was originally chemotyped as producing prymnesin type B (Binzer et al. 2019), but clustered with type A strains in our analysis. Edge length represent absolution SNP difference with the scale bar corresponding to 100,000 SNPs.

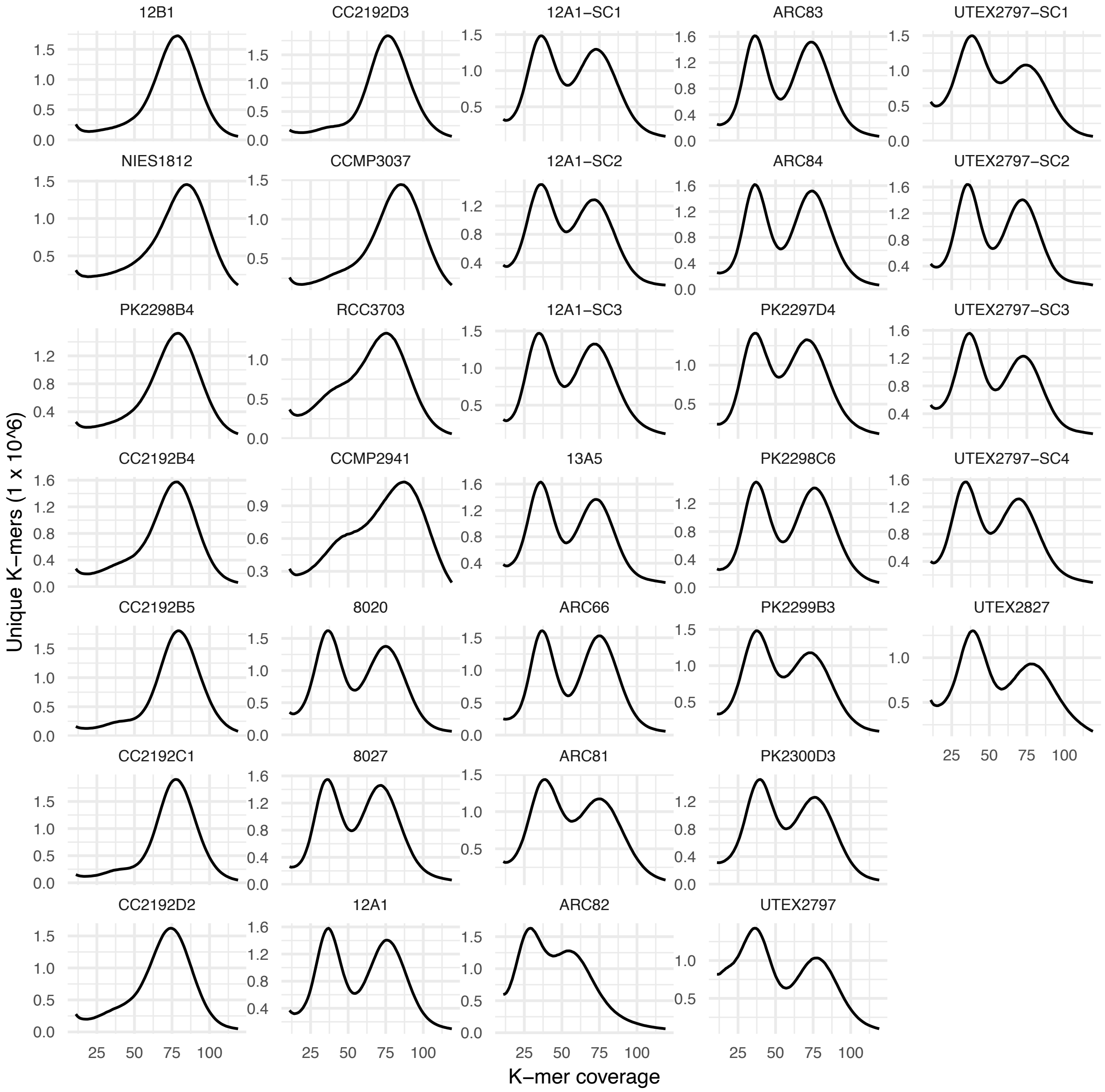


**Figure S2. Heterozygosity in *P. parvum* clade A strains.** Filtered Illumina reads were randomly subsampled to 10 million pairs per strain prior calculation of K-mer coverage to control for differences in sequencing depth between strains.

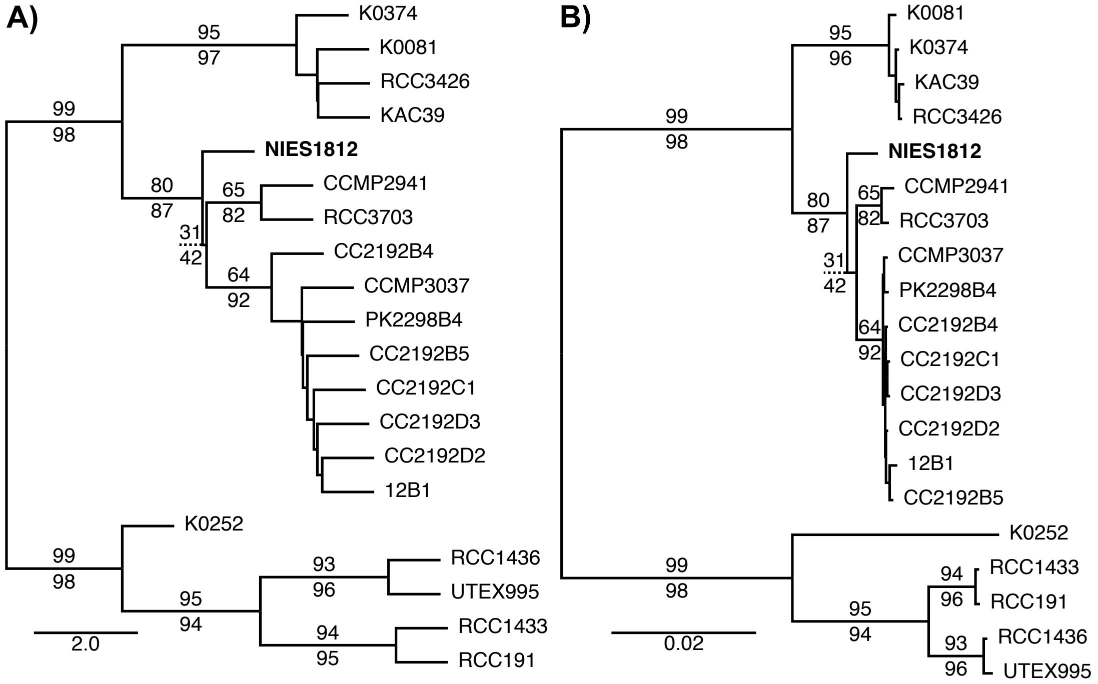


**Figure S3. Phylogenetic Trees A)** Coalescent and **B)** Concatenated species trees for the *P. parvum* s.l. complex (excluding hybrids) using 9,769 single copy genes. Support values for nodes are reported as site/gene concordance factors above/below the preceding branch. All concordance factors for major clades of *P. parvum* are reported. Ultrafast bootstrap support for all branches was ≥ 99. NIES1812 is bolded to highlight its position.

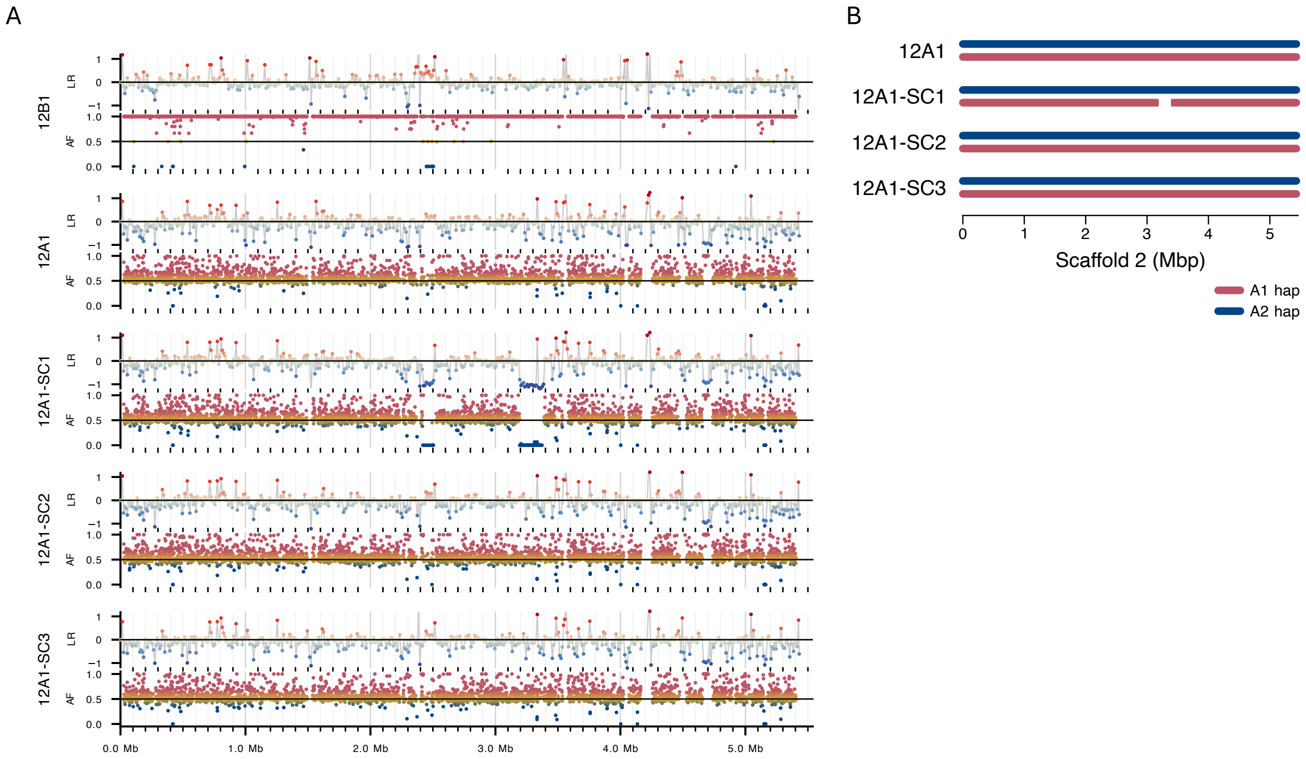


**Figure S4. Scaffold 2 structural abnormalities in 12A1 single-cell reisolates.** A) Top track indicates log2 ratio (LR) of normalized genome coverage; bottom track indicates A1 AIM allele frequency (AF; A1 read depth / total read depth). B) Model of scaffold organization.

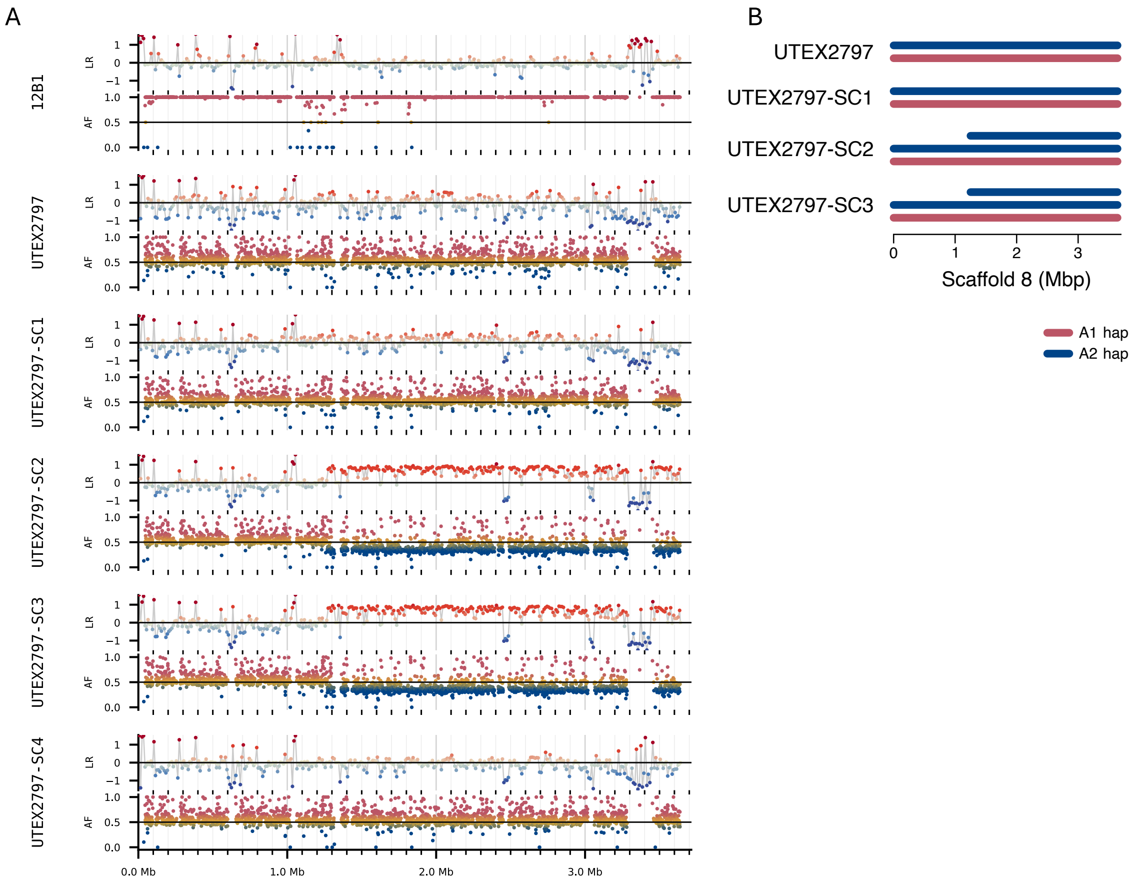


**Figure S5. Scaffold 8 structural abnormalities in UTEX2797 single-cell reisolates.** A) Top track indicates log2 ratio (LR) of normalized genome coverage; bottom track indicates A1 AIM allele frequency (AF; A1 read depth / total read depth). B) Model of scaffold organization.

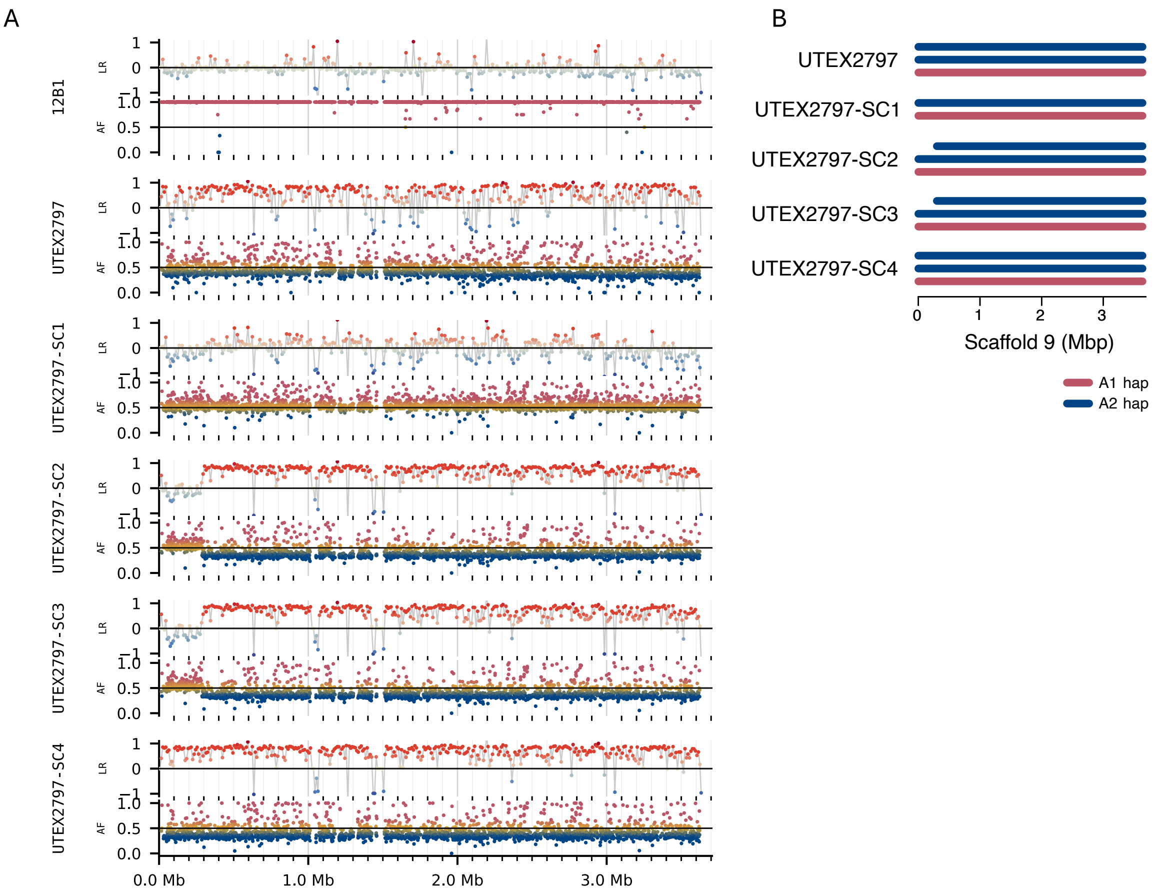


**Figure S6. Scaffold 9 structural abnormalities in UTEX2797 single-cell reisolates.** A) Top track indicates log2 ratio (LR) of normalized genome coverage; bottom track indicates A1 AIM allele frequency (AF; A1 read depth / total read depth). B) Model of scaffold organization.

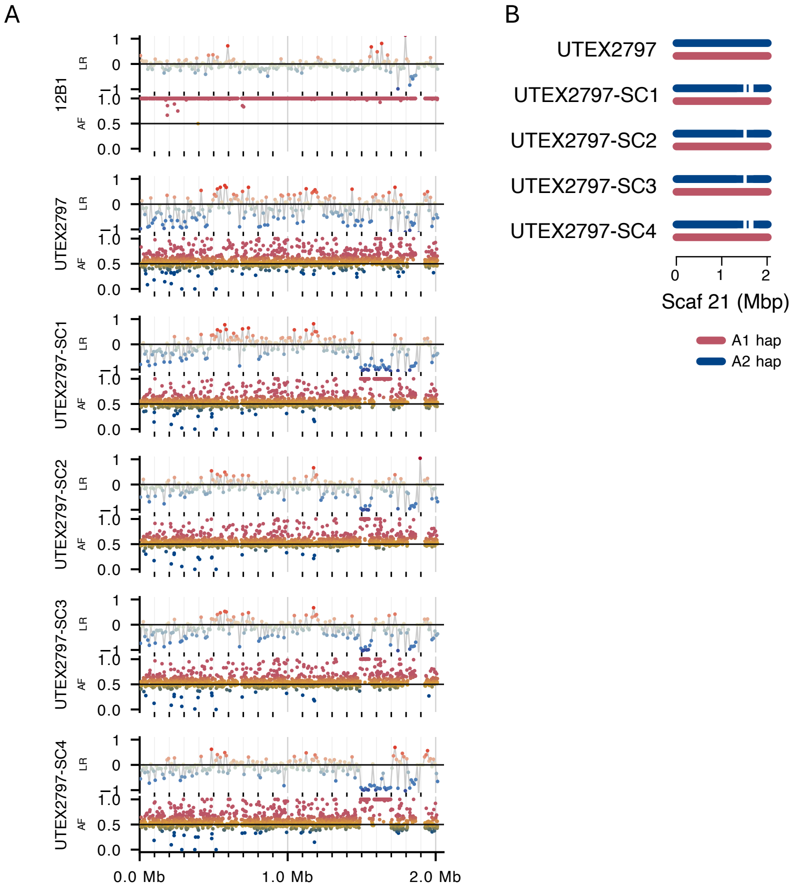


**Figure S7. Scaffold 21 structural abnormalities in UTEX2797 single-cell reisolates.** A) Top track indicates log2 ratio (LR) of normalized genome coverage; bottom track indicates A1 AIM allele frequency (AF; A1 read depth / total read depth). B) Model of scaffold organization.

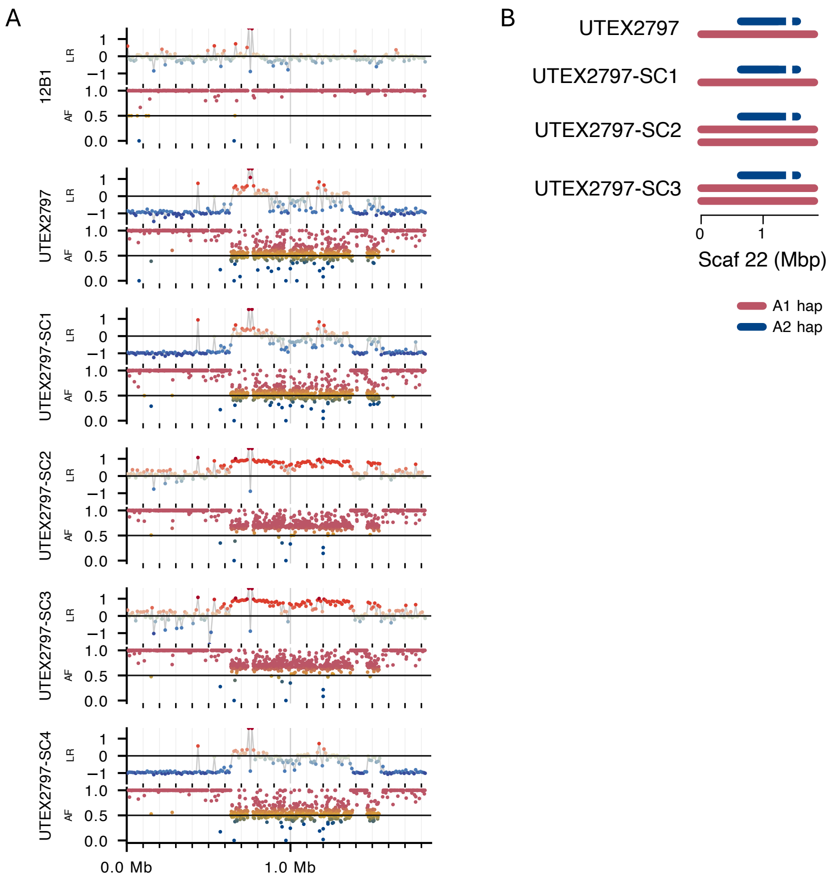


**Figure S8. Scaffold 22 structural abnormalities in UTEX2797 single-cell reisolates.** A) Top track indicates log2 ratio (LR) of normalized genome coverage; bottom track indicates A1 AIM allele frequency (AF; A1 read depth / total read depth). B) Model of scaffold organization.

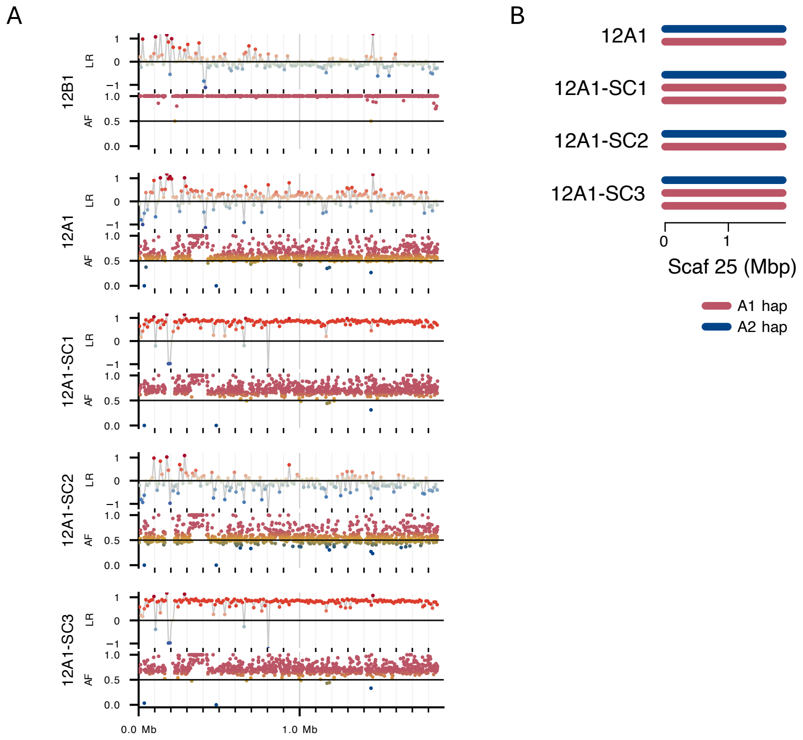


**Figure S9. Scaffold 25 structural abnormalities in 12A1 single-cell reisolates.** A) Top track indicates log2 ratio (LR) of normalized genome coverage; bottom track indicates A1 AIM allele frequency (AF; A1 read depth / total read depth). B) Model of scaffold organization.

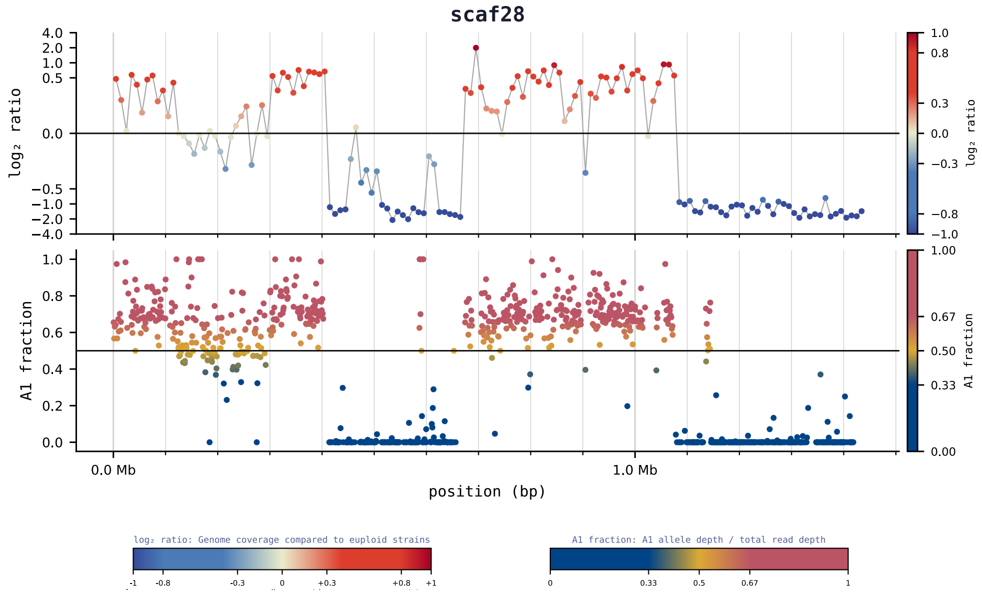


**Figure S10. Variation in coverage and allele frequency in Scaffold 28 in strain ARC82.** Top track indicates log2 ratio of normalized genome coverage; bottom track indicates A1 AIM allele frequency (A1 read depth / total read depth). Linear plots for all scaffolds for all strains are available on FigShare (see Data Availability).

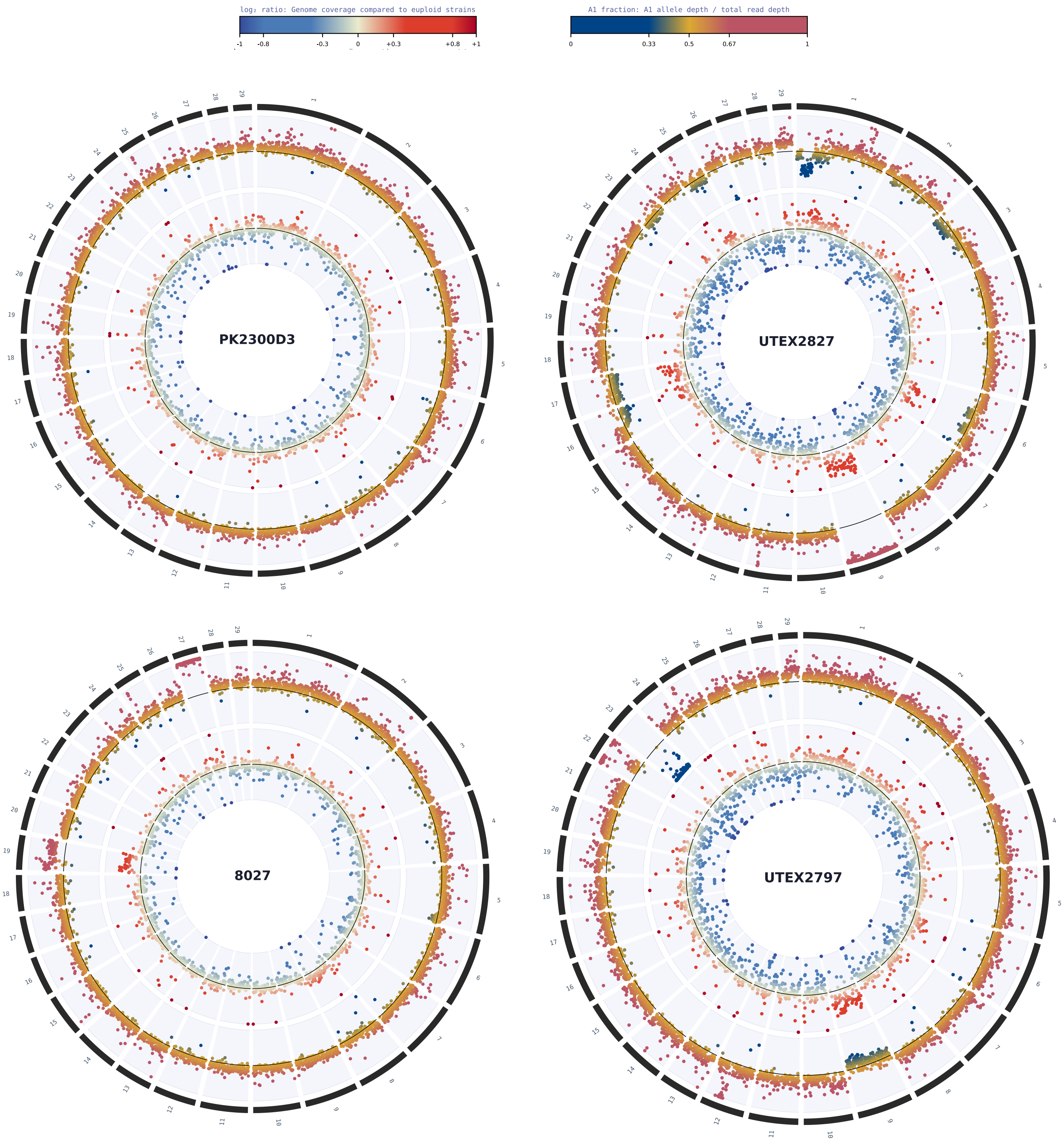


**Figure S11. Circular genome plots.** Inner tracks indicate log2 ratio of genome coverage; outer tracks indicatesA1 allele fraction (A1 read depth / total read depth). Strain PK2300D3 (top left) showed no structural anomalies. Strain UTEX2797 and 8027 (bottom plots) how examples of abnormalities on Scaffolds 9, 19, 22, 23, and 27. High variability in genome coverage and allele frequency in strain UTEX2827 (top right) prevented an accurate count of structural anomalies. Circular plots for all strains are available on FigShare (see Data Availability)

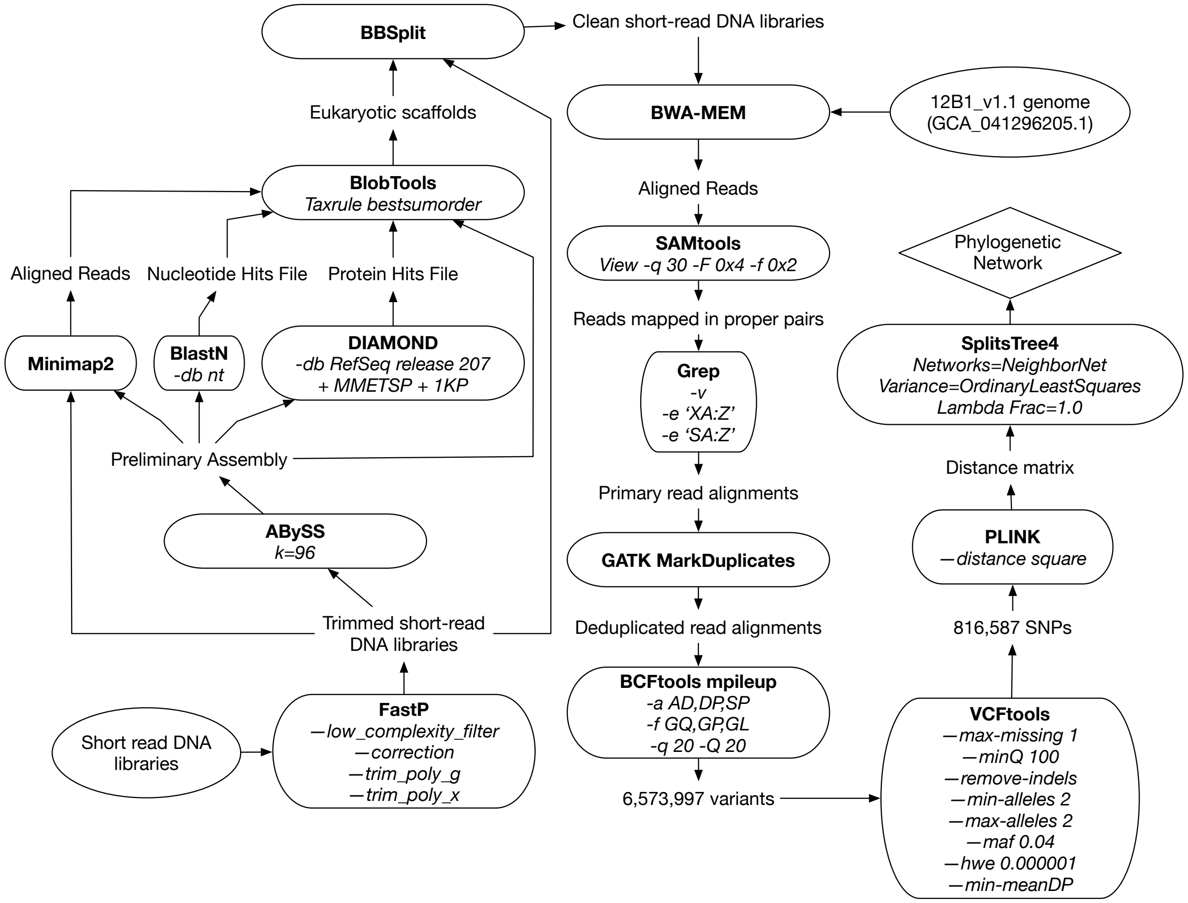


**Figure S12. Preliminary Network Flow chart.** Flow chart describing the process from short-read libraries to phylogenetic network used to identify A-type strains.

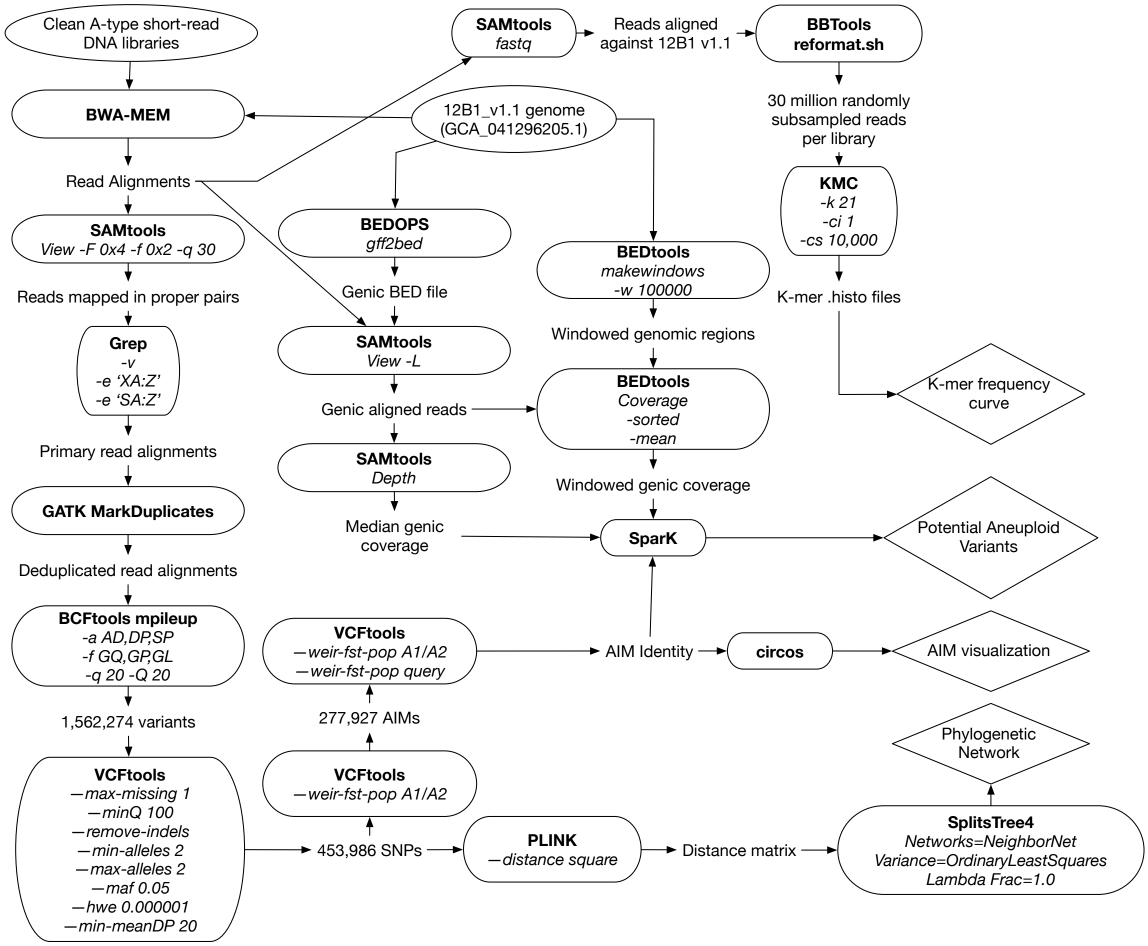


**Figure S13. A-type methods flow chart.** Flow chart describing A-type specific analyses, note that filtered short-read libraries are identical to those created at the start of **Figure S12**.

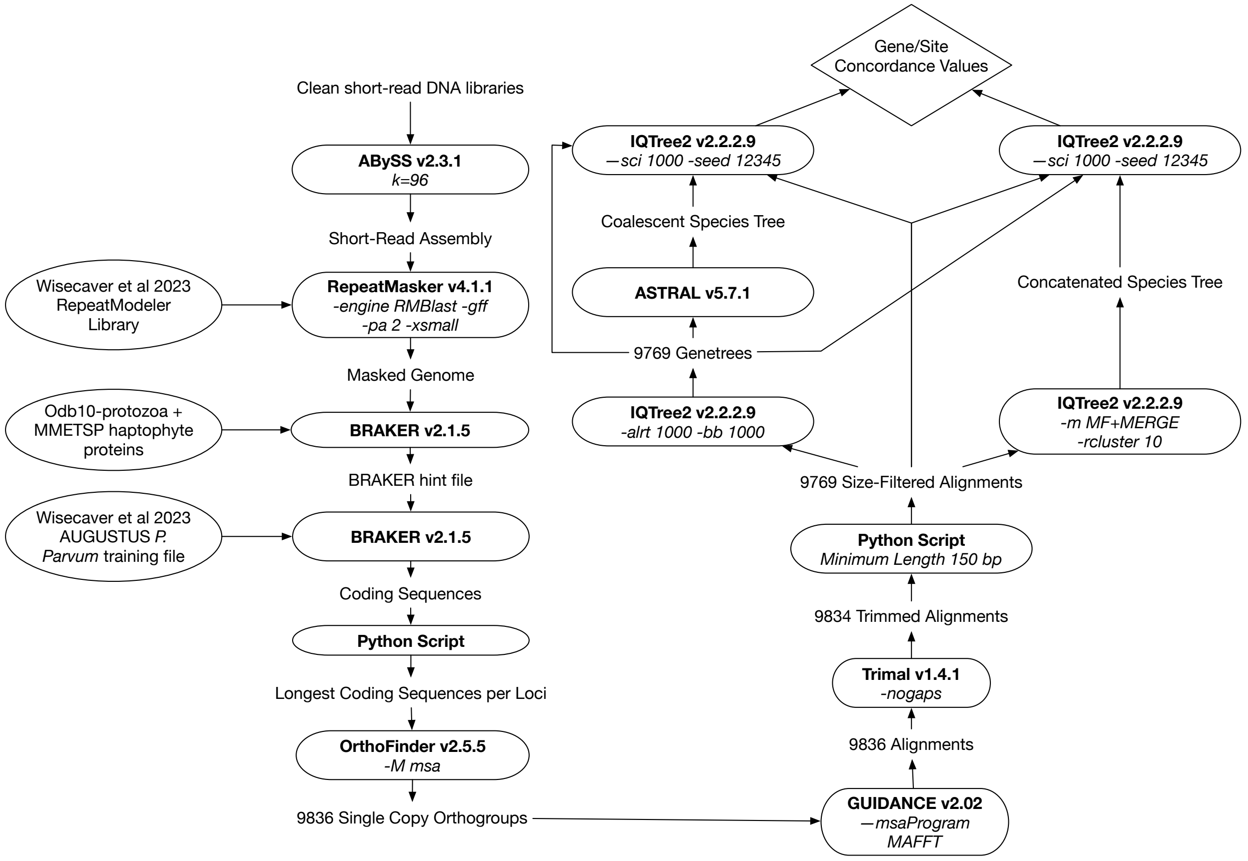


**Figure S14. Species tree creation methods flow chart.** Flow chart depicting Orthofinder analysis of all non-hybrid A, B, and C-strains, note that filtered short-read libraries are identical to those created at the start of **Figure S12.**

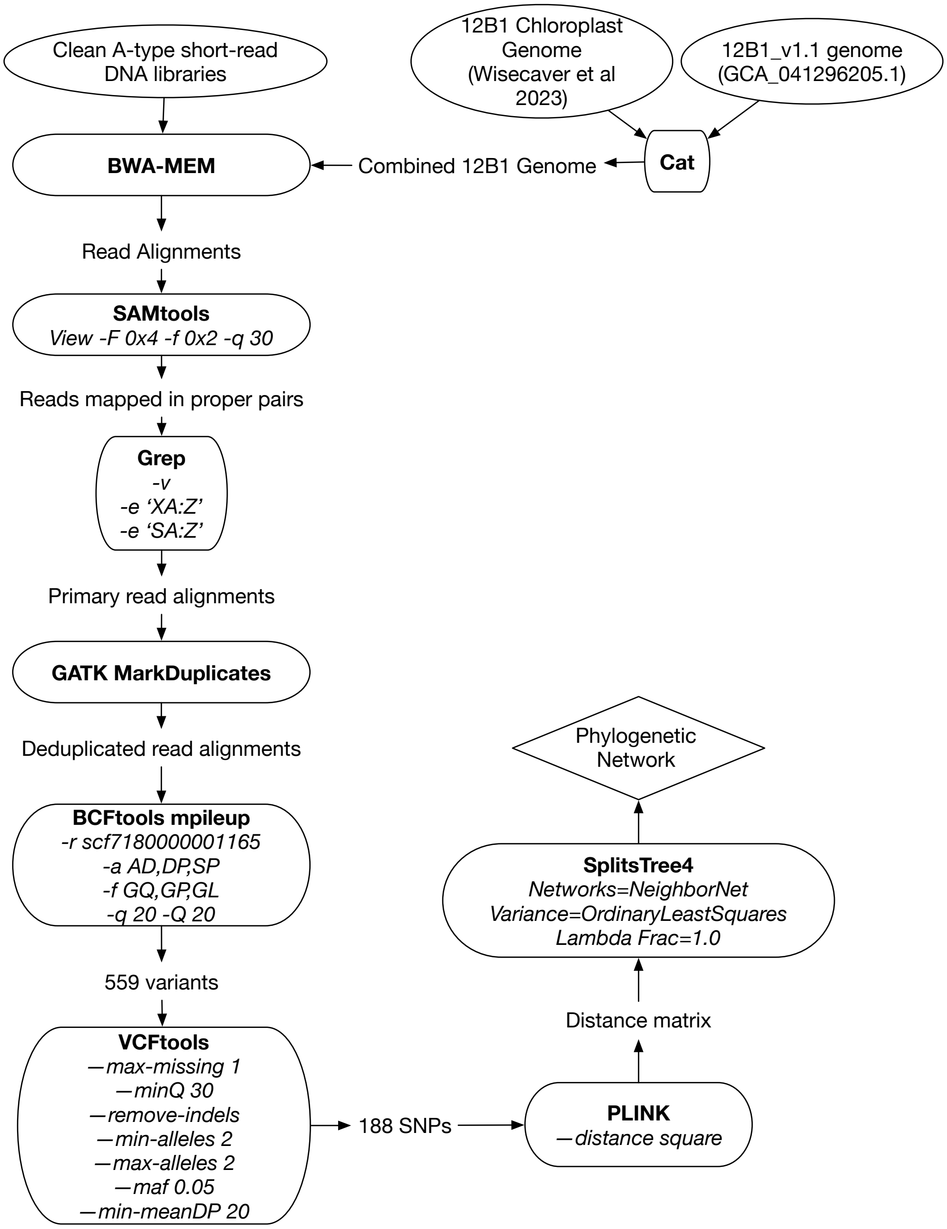


**Figure S15. Chloroplast methods flow chart.** Flow chart describing chloroplast specific analyses.
